# Limitations in PPAR⍺-dependent mitochondrial programming restrain the differentiation of human stem cell-derived β cells

**DOI:** 10.1101/2024.07.26.605318

**Authors:** Anne C. Lietzke, Emily M. Walker, Elizabeth Bealer, Kelly Crumley, Jessica King, Ava M. Stendahl, Jie Zhu, Gemma L. Pearson, Elena Levi-D’Ancona, Belle Henry-Kanarek, Rebecca K. Davidson, Jin Li, Emma C. Reck, Yifei Wu, Manikanta Arnipalli, John Paul Pham, Lakshmi Mundada, Vaibhav Sidarala, Todd J. Herron, Maria M. Coronel, Subramaniam Pennathur, Jesper G.S. Madsen, Lonnie D. Shea, Scott A. Soleimanpour

## Abstract

Pluripotent stem cell (SC)-derived islets offer hope as a renewable source for β cell replacement for type 1 diabetes (T1D), yet functional and metabolic immaturity may limit their long-term therapeutic potential. Here, we show that limitations in mitochondrial transcriptional programming impede the formation of SC-derived β (SC-β) cells. Utilizing transcriptomic profiling, assessments of chromatin accessibility, mitochondrial phenotyping, and lipidomics analyses, we observed that SC-β cells exhibit reduced oxidative and mitochondrial fatty acid metabolism compared to primary human islets that are related to limitations in key mitochondrial transcriptional networks. Surprisingly, we found that reductions in glucose-stimulated mitochondrial respiration in SC-islets were not associated with alterations in mitochondrial mass, structure, or genome integrity. In contrast, SC-islets show limited expression of targets of PPAR⍺, which regulate mitochondrial programming, yet whose functions in β cell differentiation are unknown. Importantly, treatment with WY14643, a potent PPAR⍺ agonist, induced expression of mitochondrial targets, improved insulin secretion, and increased the formation of SC-β cells both *in vitro* and following transplantation. Thus, PPAR⍺-dependent mitochondrial programming promotes the differentiation of SC-β cells and may be a promising target to improve β cell replacement efforts for T1D.

## INTRODUCTION

All forms of diabetes are associated with reduced mass and function of pancreatic islet β-cells, rendering the body unable to produce sufficient insulin to meet metabolic needs^1^. Thus, replacement of insulin-producing β cells remains a primary goal for diabetes therapy. Pluripotent stem cell (SC)-derived islets, which consist of a heterogeneous mixture of β cells, other islet hormone cell types, as well as immature progenitors and polyhormonal cells, are a promising source for β cell replacement, as these cells could be generated at scale to meet the substantial needs for patients with type 1 diabetes^2,3^ (T1D). Directed differentiation of stem cells towards the β cell lineage requires a relatively lengthy (∼1 month), complex, multi-stage protocol, including sequential use of small molecules and growth factors, cellular enrichment approaches, and modulation of spatial environments, which support the progression of SCs to progenitor cell types and eventually towards islets^4–8^. Despite the many advances in SC-islet differentiation, these protocols produce a relatively lower frequency of insulin^+^ β-like cells that are not fully functional *in vitro* and have limited glucose-stimulated insulin secretion (GSIS) compared to β cells within primary human islets.

Optimal mitochondrial function is essential to maintain GSIS, yet mitochondrial respiration is impaired in SC-islets and is of unclear etiology^5,6,9,10^. Limitations in mitochondrial metabolism of SC-islets also persist following transplantation *in vivo*^6,10,11,12^. Further, several key aspects of mitochondrial metabolism in SC-islets, such as mitochondrial fatty acid metabolism, remain unexplored. Mitochondria are vital to the function of metabolic tissues, yet recent studies position mitochondria as more than client organelles tasked with energy production and have also been shown to regulate cell fate^13,14^. However, whether mitochondrial programming can promote the SC-derived β (SC-β) cell lineage is unknown.

Here, we hypothesized and tested how targeting mitochondrial programming would improve the differentiation and function of SC-β cells. The objectives of our studies were to carefully dissect the mitochondrial metabolic function of SC-β cells, interrogate mitochondrial programming across β cell differentiation, and explore potential mitochondrial programming pathways that could be targeted to enhance mitochondrial function and, ultimately, SC-β differentiation. Together, we speculated our studies would mechanistically reveal how limitations in mitochondrial metabolism serve as an impediment to optimal SC-β cell differentiation.

## RESULTS

### SC-islets are transcriptionally and metabolically immature

We generated SC-islets via directed differentiation of the H1 human embryonic stem (hES) cell line towards a β cell fate utilizing a protocol developed by the Millman laboratory^4^ (Fig. 1A and S1A). To evaluate changes in transcriptional signatures throughout differentiation, as well as markers of islet maturity, we performed RNA-sequencing (RNA-seq) across all differentiation stages with comparison to human islet preparations from non-diabetic donors (Fig. S1B-E). These studies, as well as immunofluorescence and flow cytometry studies, demonstrated that SCs, and ultimately SC-islets, behaved largely as expected throughout differentiation, with expression of stage-specific markers and expression of islet-specific hormones in stage 6 (as well as polyhormonal cells; Fig. 1B-C and S1A). SC-islets also exhibited reduced expression of islet hormones, with exception of *GHRL*, lower levels of β and ⍺ cell maturation markers, relative hyporesponsiveness of insulin release to glucose- and KCl-stimulation, and lower insulin content compared to primary human islets (Fig. 1D-F).

**Figure 1.**
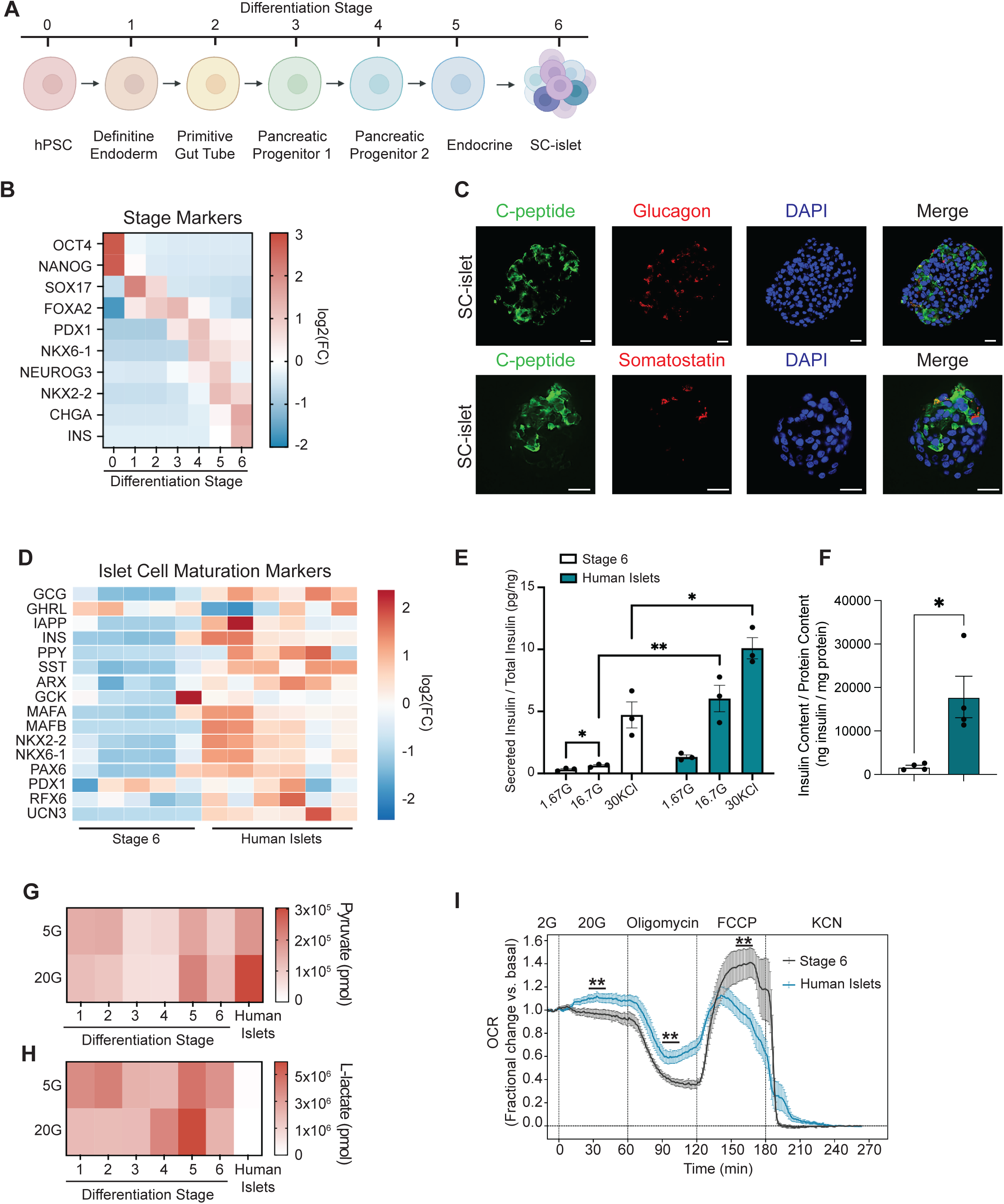
SC-islets are transcriptionally and metabolically immature. (A) A schematic diagram illustrating the cell stages during differentiation of human pluripotent stem cells (hPSCs) into SC-islets. (B) Differential RNA expression heatmap, displayed as integrated mean log_2_ fold change (FC), from RNA-seq data of representative stage marker expression at each differentiation stage (*n =* 4-5 differentiations or independent human islet donors/group). (C) Immunofluorescence images at 60X magnification of stage 6 SC-islets for C-peptide (green), Glucagon (red), Somatostatin (red), and DAPI (blue) (*n =* 3/group). Scale bars, 20 μm. (D) Differential RNA expression heatmap of genes encoding islet cell maturation markers and hormones in stage 6 SC-islets (*n =* 5) and non-diabetic human islet donors (*n =* 6). FC is relative to the average across all individual samples. (E) Insulin secretion following static incubations in 1.67 mM glucose (1.67G), 16.7 mM glucose (16.7G), and 30 mM KCl (30 KCl) normalized to insulin content in stage 6 SC-islets and human islets (*n =* 3 differentiations or independent human islet donors/group); \**P <* 0.05, \*\**P <* 0.01 by two-way ANOVA followed by Tukey’s multiple comparisons test. (F) Islet total insulin content normalized to protein content in stage 6 SC-islets and human islets (*n =* 5 differentiations and 4 independent human islet donors/group); \**P <* 0.05 unpaired *t*-test. Differential metabolomics heatmap demonstrating mean content of pyruvate (G) or lactate (H) at 5 mM glucose (5G) and 20 mM glucose (20G) for differentiation stages 1 through 6 (*n =* 3/group) and human islets (*n =* 5 independent islet donors). (I) Oxygen consumption rate (OCR) fractional change in stage 6 SC-islets (*n =* 4 differentiations) and human islets (*n =* 5 independent islet donors) following exposure to 2 mM glucose (2G), 20 mM glucose (20G), 10 μM oligomycin, 1 μM FCCP, and 3 mM KCN. Data are presented as mean from independent differentiations or donors ± SEM; \**P <* 0.05, \*\**P <* 0.01 by unpaired *t*-test of fractional OCR value at designated time point for minimal or maximal respiration by oligomycin or FCCP, respectively.

To complement our findings of transcriptional and functional immaturity in SC-islets, we next profiled metabolic endpoints in SC-islets. We first surveyed the glycolytic intermediates pyruvate and lactate by LC-MS, given the high importance of pyruvate utilization in mature β cells^15–18^. We found that stage 5 and stage 6 cells increased pyruvate concentrations in response to glucose stimulation, albeit at lower levels than human islets (Fig. 1G). Moreover, SCs at every stage generated significantly higher quantities of lactate, including in stage 6 cells, as compared to human islets (Fig. 1H). The presence of higher quantities of lactate in SC-islets than human islets could be consistent with previous observations of defects in glycolytic flux, an inability to silence lactate metabolism due to expression of disallowed genes (such as *LDHA*), or due to the inability to switch from primarily glycolytic to oxidative metabolism in immature β cells, all of which have been reported previously^5,6,10,18,19^. We next performed respirometry studies utilizing the recently described Barofuse instrument, which allows for the assessment of respiration in a gas pressure flow-based perifusion style platform^20^. We confirmed that SC-islets did not increase oxygen consumption upon glucose stimulation as compared to human islets, which has been shown previously^5,6,10^ (Fig. 1I). We also observed that SC-islets displayed more sensitivity to the effects of the ATP synthase inhibitor oligomycin, which is expected to lower oxygen consumption, and the uncoupler FCCP, which transiently increases oxygen consumption, than primary human islets, possibly suggesting that SC-islets may not be as uncoupled as primary human islets but could also be related to relative differences in penetration of the compounds (Fig. 1I). Taken together, SC-islets display a signature of transcriptional, functional, and metabolic immaturity.

### Mitochondrial DNA content, mass, and architecture in SC-islets are comparable to those of human islets

Optimal β cell function is highly dependent on mitochondrial quality and health^21–23^. Given our observations of transcriptional, functional, and metabolic immaturity in SC-islets, we profiled mitochondrial mass and structure of SC-islets at different stages of differentiation and human islets to obtain insights into limitations in mitochondrial respiration as recent studies have had discordant results regarding reductions in mitochondrial mass in SC-islets compared to primary human islets^5,10^. To survey mitochondrial genome integrity, we measured mitochondrial (mt)DNA content, and while we found higher mtDNA content in stage 0 embryonic stem cells, we noted no significant differences between stage 1-6 cells and human islets (Fig. 2A). To next determine if there were deficiencies in mtDNA replication, we evaluated the initiation of first strand replication. In mtDNA replication, first strand synthesis begins with formation of the D-loop. Elongation of the 7S sequence, which is contained in the D-loop, into the nearby region, *Cytb*, signifies committed initiation of first strand replication. We assessed both D-loop formation and first strand replication initiation and found no differences between stage 0-6 cells and human islets (Fig. 2B). We also observed no differences in mitochondrial mass in stage 0-6 cells compared to human islets by use of two complementary approaches; namely, citrate synthase activity and TOM20 protein expression (Fig. 2C-D and S1F). To assess mitochondrial ultrastructure, we next performed transmission electron microscopy. While we noted a lower frequency of mature insulin granules, as previously described^5,6,24,25^, we did not observe any differences in quantitative measures of mitochondrial architecture in stage 6 SC-β cells compared to primary human β cells (Fig. 2E). Together, these results suggest that limitations in mitochondrial respiration in SC-islets were not a consequence of reduced mitochondrial genome integrity, DNA replication, mass, or structure.

**Figure 2.**
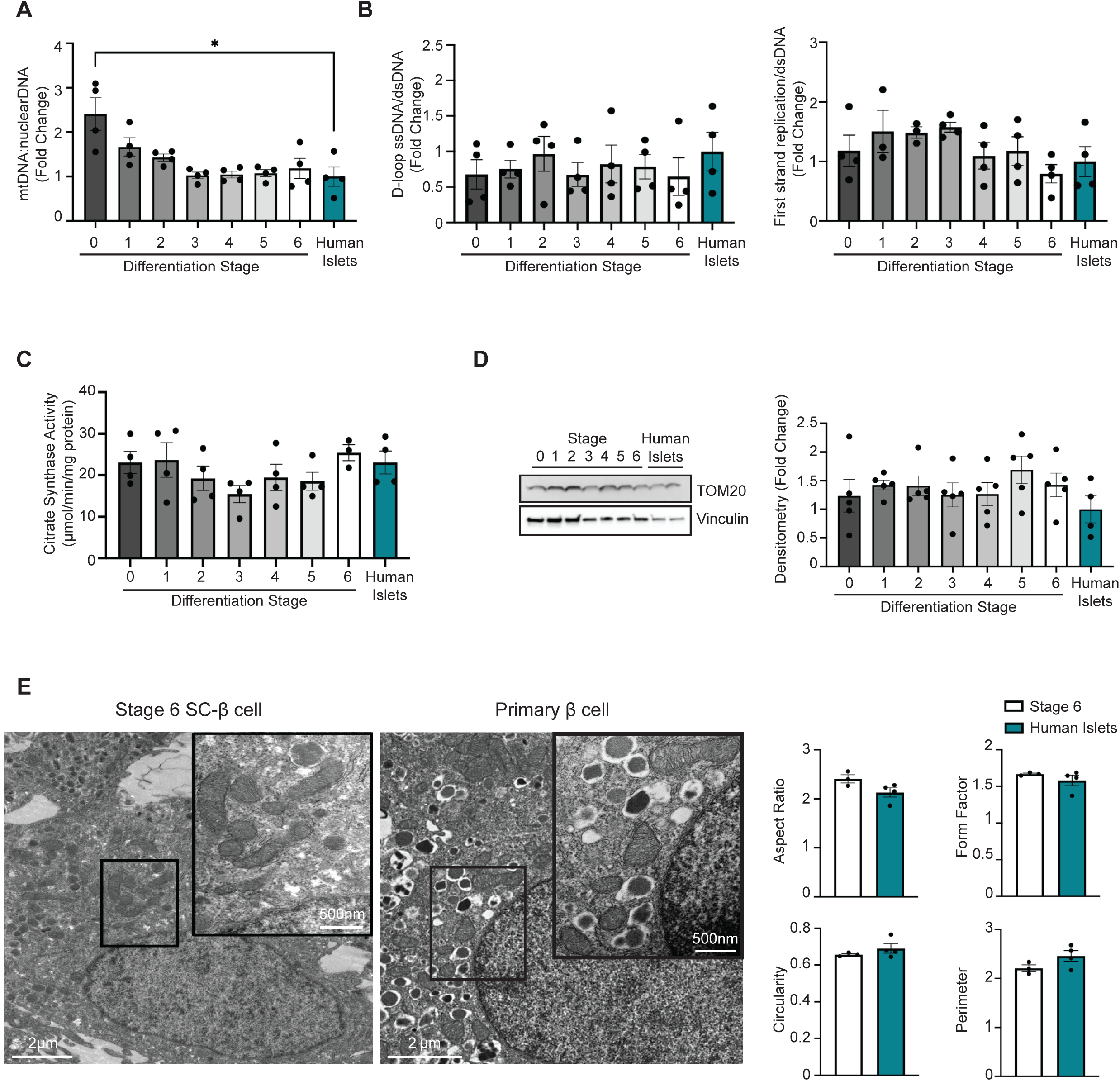
SC-islets display comparable mitochondrial DNA content, mass, and architecture to human islets. (A) Quantification of ratio of mitochondrial DNA to nuclear DNA by qPCR for stage 0-6 cells and human islets (*n =* 4/group). (B) Quantification of single strand mitochondrial DNA products measured by qPCR for stage 0-6 cells and human islets (*n =* 3-4/group). (C) Quantification of citrate synthase activity normalized to total protein content for stage 0-6 cells and human islets (*n =* 3-4/group). (D) Representative Western blot (WB) of TOM20 protein expression for stage 0-6 cells and two different human islet donors (left). Vinculin serves as a loading control. Quantification (right) of TOM20 expression (normalized to Vinculin expression) via densitometry of the Western blots (*n =* 4-5/group). (E) Representative transmission electron microscopy (TEM) images of stage 6 SC-β cells and primary human β cells (left) and quantification of mitochondria ultrastructure (right) (*n =* 3/group, ∼150 β cell mitochondria scored/sample).

### Transcriptomic profiling reveals potential limitations in mitochondrial metabolism in SC-islets

Due to the similarities in mitochondrial genome integrity, DNA replication, mass, and structure between SC-islets and human islets, we next explored potential transcriptional differences that might provide mechanistic insights into limitations in glucose-stimulated mitochondrial respiration in SC-islets. Following a comparison of RNA-seq data between stage 6 SC-islets and human islets, we found that 11,858 genes were significantly differentially expressed, including 405 genes represented within MitoCarta 3.0^26^, a curated set of nuclear and mitochondrial-encoded genes with high confidence of localization to the mitochondria (Fig. 3A). While we observed dynamic changes in expression of MitoCarta genes throughout differentiation, we noted a subset of highly expressed MitoCarta genes in primary human islets that appeared to remain relatively lowly expressed throughout SC differentiation, including by stage 6 (Fig. 3B). Pathways analysis of differentially expressed MitoCarta genes showed differences in fatty acid metabolism, amino acid metabolism, and oxidative phosphorylation (OXPHOS) (Fig. 3C-D). As there were a greater number of downregulated Mitocarta 3.0 genes in SC-islets compared to human islets, we wanted to understand the possible biological processes associated with these gene targets via the Search Tool for Recurring Instances of Neighboring Genes (STRING) database^27^. Following STRING analysis of significantly downregulated MitoCarta genes from stage 6 SC-islets, gene ontology (GO) enrichment analysis was used to clarify pathways related to connections identified by STRING. Pathways identified were related to mitochondrial translation, the electron transport chain (ETC)-OXPHOS system, and lipid metabolism (Fig. 3E). Collectively, our transcriptomic studies revealed several pathways that could contribute to limitations in mitochondrial metabolism in SC-islets.

**Figure 3.**
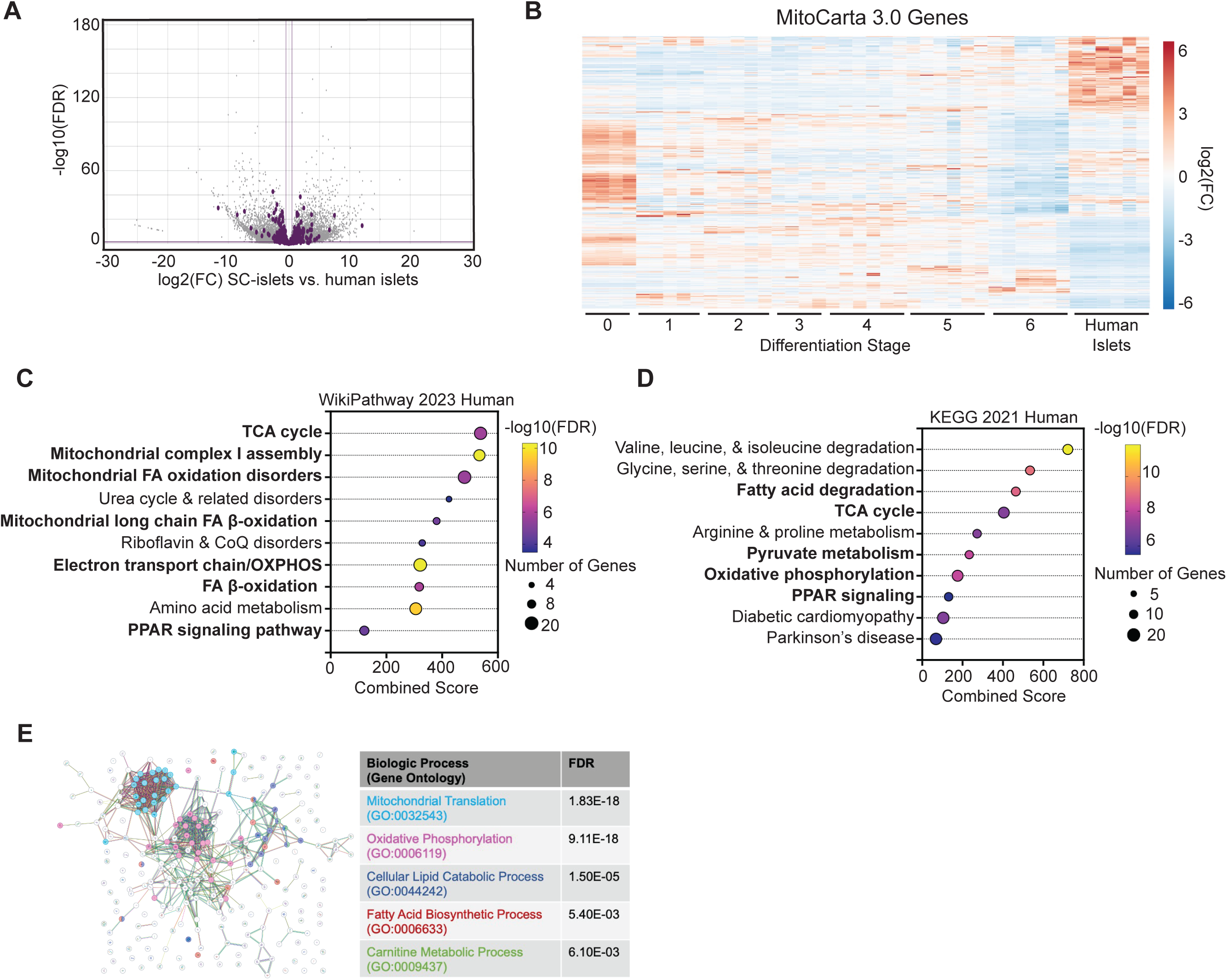
Transcriptomic profiling of SC-islets versus human islets reveals potential limitations in mitochondrial metabolism. (A) Volcano plot depicting differential RNA expression in stage 6 SC-islets compared to human islets (*n =* 6 differentiations or independent human islet donors/group). Significantly differentially expressed genes demarcated by -FDR > 0.5 and |FC| > 1.4. MitoCarta 3.0 genes are highlighted in purple. (B) Differential RNA expression heatmaps from RNA-seq data demonstrating expression of significantly differentially expressed MitoCarta 3.0 genes in stage 0-6 cells compared to human islets (*n =* 4-6 differentiations or independent human islet donors/group). FC is relative to the average across all individual samples. Selected terms from Enrichr analysis of differentially expressed MitoCarta genes from stage 6 SC-islets vs. human islets analyzed with the following libraries: (C) WikiPathway 2023 Human and (D) KEGG 2021 Human. (E) STRING analysis of MitoCarta genes with reduced expression in stage 6 SC-islets compared to human islets.

### Limitations in the OXPHOS machinery and mitochondrial lipid metabolism in SC-islets

We next wished to clarify the specific nature of transcriptional defects affecting mitochondria in SC-islets. An initial question was whether the limitations in SC-islet mitochondria were related to an impairment in the shift of the glycolytic to OXPHOS metabolic program, which was previously observed in human induced pluripotent stem cell (iPSC)-derived islets^11^. While we observed that expression of genes related to glucose transport and glycolysis were enriched in stage 0 pluripotent cells, we did not observe differences in the glycolytic gene expression program between stage 6 SC-islets and human islets (Fig. 4A). We also did not observe a significant difference in the overall RNA expression of genes encoding the entirety of TCA cycle enzymes or enzymes/transporters involved in lactate metabolism between stage 6 SC-islets and human islets, whereas these genes were more highly enriched in stage 0 ES cells (Fig. S1G). We next considered the possibility that mitochondrial impairments in SC-islets could be related to limitations in the β cell OXPHOS machinery. Indeed, we observed a notable subset of 29 OXPHOS gene targets that were enriched in human islets compared to stage 6 SC-islets (Fig. 4B-C). These upregulated targets in human islets included genes encoding subunits of all 5 ETC complexes, as well as complex assembly factors (Fig. 4B-C). Notably, stage 0 pluripotent cells also had an enrichment of a subset of OXPHOS gene targets compared to stage 1-6 cells; however, these genes appeared to be largely distinct from those upregulated within human islets (Fig. 4B). To confirm that this phenotype was specific to SC-β cells compared to other SC-islet cell types, we re-analyzed data from a recently published single-cell RNA-seq dataset comparing SC-islets and human islets, as it has been observed that the various directed differentiation protocols from independent laboratories produce SC-islets with similar transcriptomic profiles^28,29^. Indeed, we observed that >50% of OXPHOS gene targets were lower in SC-β cells compared to primary human β cells, whereas there was no significant differences in glycolytic gene expression between the groups (Fig. 4D). Reduced expression of OXPHOS genes appeared to be β cell-specific, as we did not find robust differences in SC-⍺ cells compared to primary human ⍺ cells (Fig. 4D). Lower expression of ETC and OXPHOS gene targets in SC-β cells was also supported by transcriptomic profiling by others^6^. We next measured expression of subunits of OXPHOS complexes via Western blot and found that their expression was significantly lower in SCs compared to human islets across all differentiation stages (Fig. 4E). Thus, we observed lower expression of components of the OXPHOS machinery in SCs compared to human islets that appear to be distinct from programming of the glycolytic gene network.

**Figure 4.**
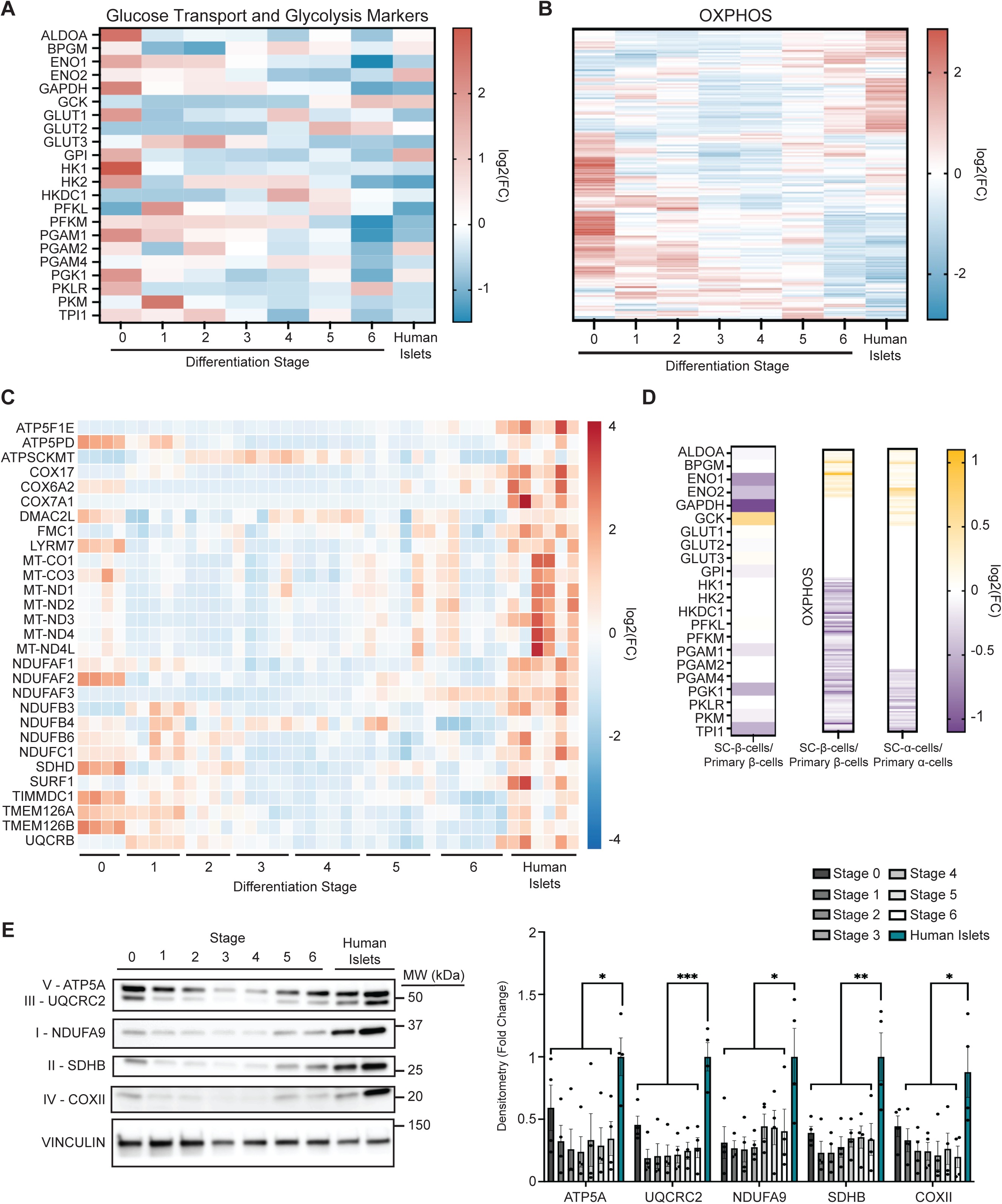
Reduced expression of the OXPHOS machinery in SC-islets. (A) Differential RNA expression heatmap, displayed as integrated mean log_2_ fold change (FC), from RNA-seq data of genes related to glucose transport and glycolysis at each differentiation stage and human islets (*n =* 4-5 differentiations or independent human islet donors/group). (B) Differential RNA expression heatmap (presented as integrated mean/group) of genes encoding OXPHOS components at each differentiation stage as well as human islet donors (*n =* 4-6/group). (C) Differential RNA expression heatmaps from RNAseq demonstrating expression of OXPHOS machinery genes in stage 0-6 cells and human islets (*n =* 4-6 differentiations or independent human islet donors/group). FC is relative to the average across all individual samples. (D) Differential RNA expression heatmap, displayed as integrated mean log_2_FC, from scRNA-seq of genes related to glycolysis and glucose transport and OXPHOS between SC-β cells to primary β cells as well as OXPHOS genes between SC-⍺ cells to primary ⍺ cells. Data curated from Augsornworawat *et al*.^28^. (E) Representative Western blot analysis of expression of OXPHOS subunits, including ATP5A (Complex V), UQCRC2 (Complex III), NDUFA9 (Complex I), SDHB (Complex II), and COXII (Complex IV) in stage 0-6 cells and human islets (left). Vinculin serves as a loading control. Quantification (right) of ATP5A, UQCRC2, NDUFA9, SDHB, and COXII protein expression by densitometry (normalized for Vinculin expression) of the Western blots of stage 0-6 cells and human islets (*n =* 4 differentiations or independent human islet donors/group); \**P <* 0.05, \*\**P <* 0.01, \*\*\**P <* 0.001, by one-way ANOVA followed by Dunnett’s multiple comparisons test.

While β cells are not classically considered central regulators of physiologic lipid storage and turnover, lipid metabolism and signaling are well-known to be essential for the maintenance of optimal β cell insulin release^30,31^. Importantly, mitochondrial lipid metabolism has not been examined in SC-islets to date. We observed distinct subsets of genes related to fatty acid biosynthesis and β-oxidation that were enriched in human islets but were not increased during differentiation towards stage 6 SC-islets (Fig. 5A-B). Specifically, we found genes encoding key initial steps in fatty acid biosynthesis and β-oxidation were markedly lower in SC-islets compared to human islets, including *ACACB*^32,33^, a key component of acetyl CoA carboxylase, which catalyzes conversion of acetyl CoA to malonyl CoA as an early step in fatty acid biosynthesis, *CPT1A*^32,34^, the rate-limiting enzyme of fatty acid oxidation that catalyzes the transfer of the long-chain acyl-CoA ester to carnitine, to allow fatty acids to enter the mitochondrial matrix for oxidation, and *ACAT1*^34,35^, which catalyzes the final step of mitochondrial β-oxidation (Fig. 5C).

**Figure 5.**
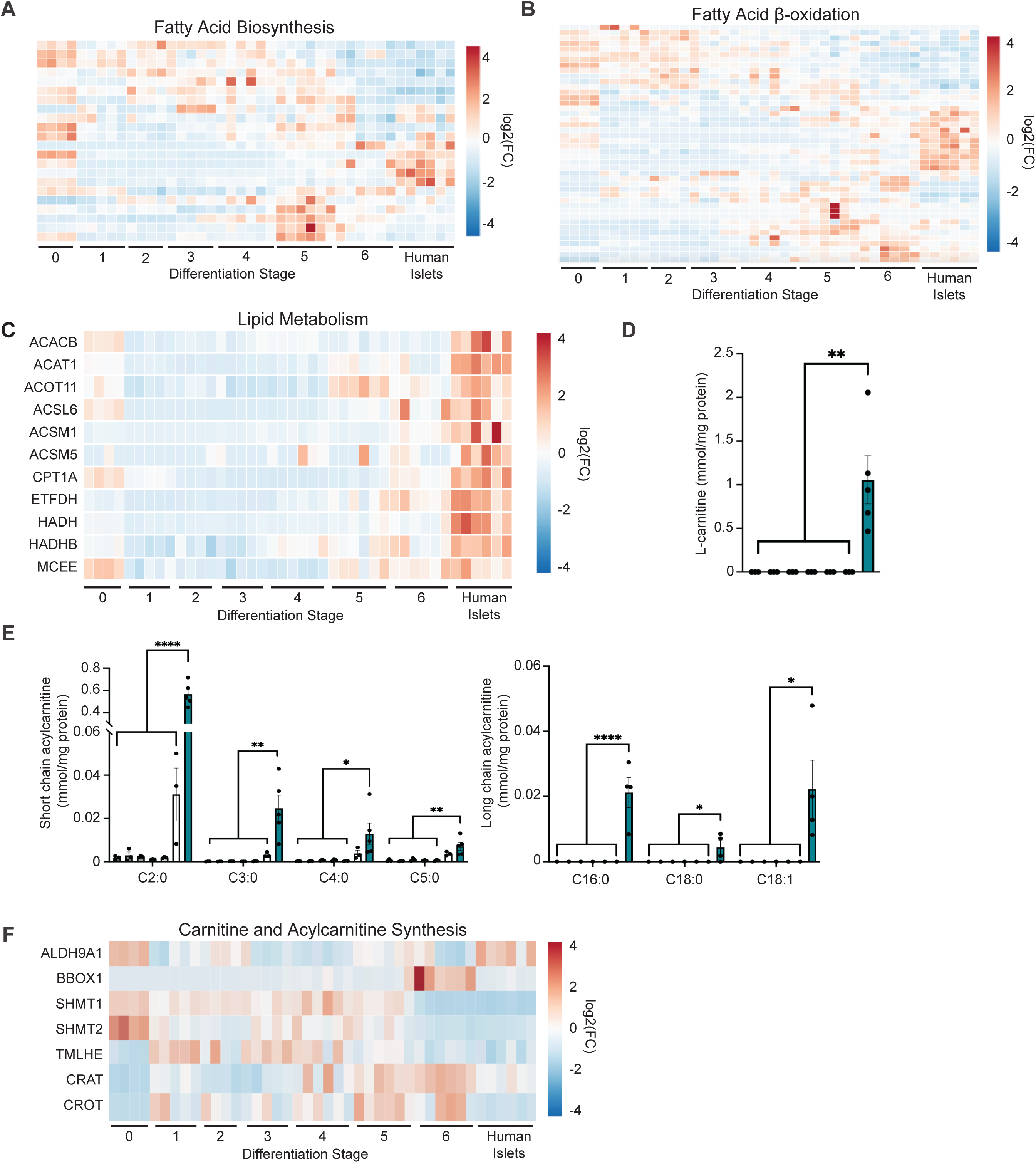
Limitations in mitochondrial lipid metabolism in SC-islets. Differential RNA expression heatmaps from RNA-seq data demonstrating expression of fatty acid biosynthesis (A) or fatty acid β-oxidation (B) genes in stage 0-6 cells and human islets (*n =* 4-6 differentiations or independent human islet donors/group). FC is relative to the average across all individual samples. (C) Differential RNA expression heatmaps from RNA-seq data demonstrating expression of lipid metabolism genes in stage 0-6 cells and human islets (*n =* 3-5 differentiations or independent human islet donors/group). FC is relative to the average across all individual samples. Lipidomics profiling of (D) L-carnitine (E) and short- or long-chain acylcarnitines (normalized to protein content) for stage 1-6 cells (*n =* 3/group) and human islets (*n =* 4-5 independent islet donors); \**P <* 0.05, \*\**P <* 0.01, \*\*\*\**P <* 0.0001 by one-way ANOVA followed by Dunnett’s multiple comparisons test. (F) Differential RNA expression heatmaps from RNA-seq data demonstrating expression of expression of carnitine biosynthesis and short- and medium-chain acylcarnitine synthesis genes in stage 0-6 cells and human islets (*n =* 3-5 differentiations or independent human islet donors/group). FC is relative to the average across all individual samples.

To assess if differentiating SCs and SC-islets possessed alterations in mitochondrial fatty acid metabolism, we examined acylcarnitine species by LC-MS, as carnitine is a required cofactor for lipid transport into the mitochondria for fatty acid β-oxidation^36^. Notably, we could not detect long-chain acylcarnitines or L-carnitine during differentiation, while these were detectable in human islets (Fig. 5D-E). We next noted low levels of short-chain acylcarnitines throughout SC differentiation, with elevations in short-chain acylcarnitines by stage 6. However, both short- and long-chain acylcarnitines were significantly reduced throughout differentiation to SC-islets when compared to human islets (Fig. 5E). Furthermore, we observed lower expression of *ALDH9A1*, a gene encoding a carnitine biosynthetic enzyme, as well as in *CPT1A*, the key regulator of long-chain acylcarnitine synthesis, in SC-islets compared to human islets, yet other regulators of short-chain acylcarnitine synthesis and other genes in the carnitine biosynthetic pathway were not lower in SC-islets (Fig. 5C and 5F). Of note, L-carnitine is not included within either SC-islet^4^ or human islet media formulations, thus these differences were unrelated to L-carnitine supplementation (B. Haight, Prodo Laboratories, personal communication as human islet media formulation is proprietary). These data indicate that in addition to limitations in expression of the ETC-OXPHOS system, SC-islets exhibit robust differences in mitochondrial fatty acid metabolism compared to primary human islets.

To ensure our transcriptomic findings were not limited to differentiation of a single hES line, we performed RNA-seq in stage 0-6 cells generated from the HUES8 hES cell line and compared these results to primary human islets. We again observed the expected expression of stage-specific differentiation markers and limitations of islet hormones and of β and ⍺ cell maturation markers in stage 6 cells, similar to our observations in H1 cells (Fig. S2A-C). Further, we observed lower expression of key regulators related to fatty acid β-oxidation, glucose transport and glycolysis genes, and genes encoding subunits of all 5 ETC complexes, complex assembly factors in stage 6 SC-islets compared to primary human islets (Fig. S2D-F). In sum, these data indicate that SC-islets have several limitations in mitochondrial programming, including within the ETC-OXPHOS system and mitochondrial fatty acid metabolism.

### SC-islets display lower expression of PPAR⍺ and PPARγ transcriptional targets

Due to the transcriptional differences in the OXPHOS machinery and mitochondrial lipid metabolism, we hypothesized that limitations in key transcriptional pathways might prevent optimal mitochondrial programming in SC-islets. To identify candidate transcriptional regulators, we employed the bioinformatic algorithm Enrichr^37–39^ to analyze differentially expressed Mitocarta 3.0 genes between stage 6 SC-islets and human islets (Fig. 6A-B). This analysis yielded several expected findings, including the estrogen-related receptor (ERR) family, PDX1, and PRDM16, which all have been associated with the regulation of β-cell development, as well as several effects on mitochondria, including mitochondrial biogenesis, mitophagy, or induction of oxidative metabolism^11,40–43^. We were also intrigued by the enrichment of PPAR⍺ and PPARγ in SC-islets, nuclear receptors that are both tightly connected with mitochondrial function, including the regulation of fatty acid oxidation, mitochondrial biogenesis, and coordination of mitochondrial programming together with the co-activator PGC1⍺, a central regulator of mitochondrial metabolism^44–46^. Thus, we next assessed the expression of PPAR⍺ and PPARγ target genes in stage 6 SC-islets and human islets as PPAR⍺ and PPARγ had not been previously studied during SC-β cell differentiation. We found that expression of canonical PPAR⍺ and PPARγ target genes were lower in stage 6 SC-islets compared to human islets (Fig. 6C). Further, PPAR⍺ regulates carnitine biosynthesis and transcriptionally regulates expression of *ALDH9A1*^47,48^, which is also lower in SC-islets compared to human islets, as noted above (Fig. 5F).

**Figure 6.**
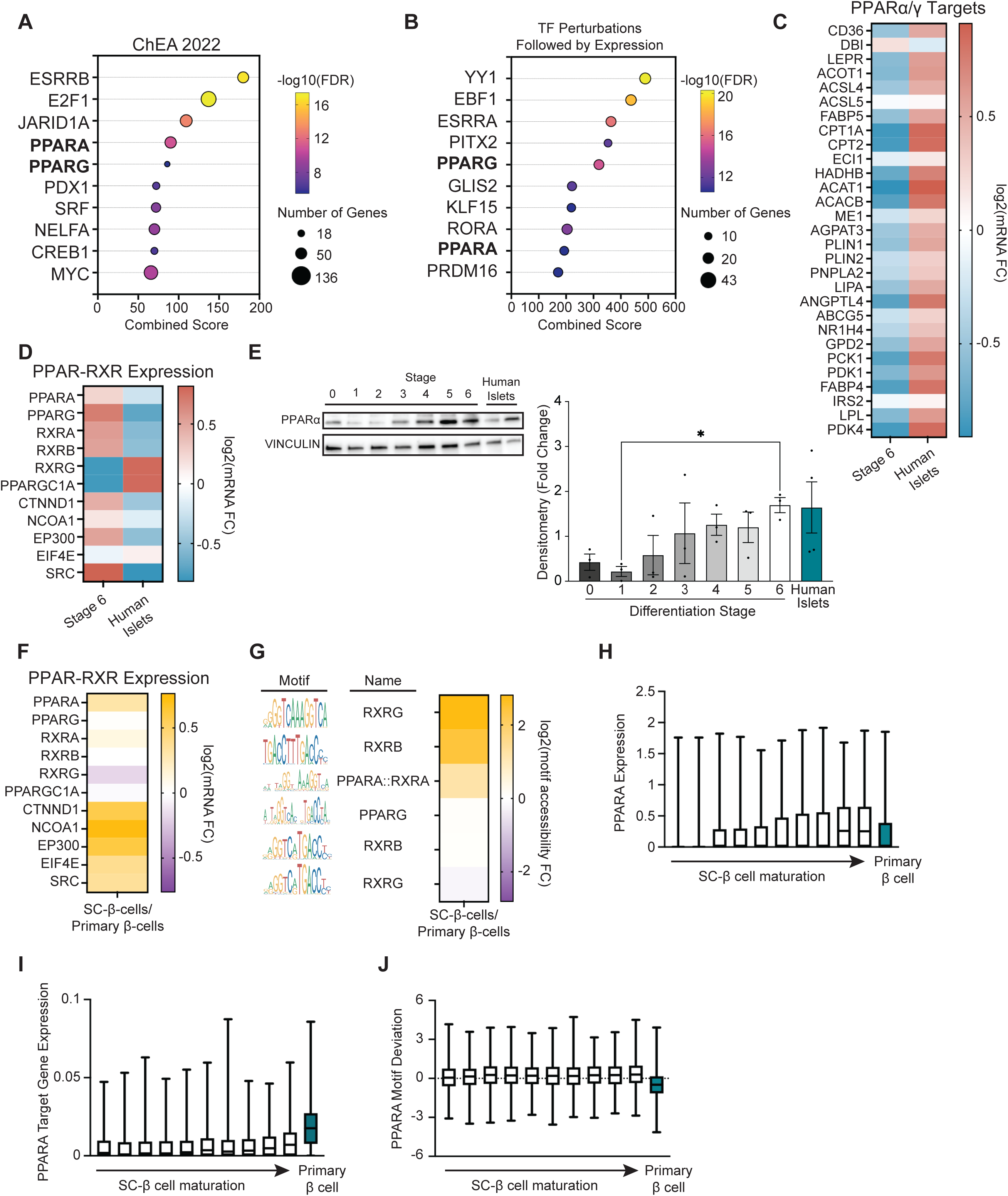
SC-islets display reduced expression of PPAR⍺ and PPARγ transcriptional targets. Enrichr analysis of significantly differentially expressed genes from stage 6 SC-islets compared to human islets analyzed with the following libraries: (A) ChEA 2022 and (B) Transcription factor (TF) perturbations followed by expression. (C) Differential RNA expression heatmap from RNA-seq data demonstrating integrated mean expression of PPAR⍺ targets for stage 6 SC-islets and human islets (*n =* 6/group). (D) Differential RNA expression heatmap from RNA-seq data demonstrating integrated mean expression of genes encoding components of the PPAR-RXR transcriptional binding complex, including transcriptional co-activators, in stage 6 SC-islets and human islets (*n =* 6/group). (E) Representative Western blot analysis of expression of PPAR⍺ in stage 0-6 cells and human islets (left). Vinculin serves as a loading control. Quantification (right) of PPAR⍺ protein expression by densitometry (normalized to Vinculin expression) of the Western blots of stage 0-6 cells and human islets (*n =* 3 differentiations or 4 independent human islet donors/group); \**P <* 0.05 by one-way ANOVA followed by Dunnett’s multiple comparisons test. (F) Differential RNA expression heatmap generated from publicly-available scRNA-seq data^28^ demonstrating integrated mean expression of genes encoding components of the PPAR-RXR transcriptional binding complex, including transcriptional co-activators, in SC-β cells and primary β cells. (G) Differential chromatin accessibility heatmap generated from publicly-available scATACseq data^28^ demonstrating integrated mean motif accessibility of PPAR and RXR binding sites in SC-β cells and primary β cells. (H) Minimum to maximum box and whisker plots of normalized expression of *PPARA* across SC-β cells differentiation trajectory and primary β cells generated from publicly-available scRNA-seq data^28^ (*n =* 3-4 differentiations or independent human islet donors/group). (I) Minimum to maximum box and whisker plots of PPARA module scores across SC-β cells differentiation trajectory and primary β cells generated from publicly-available scRNA-seq data^28^ representing PPARA target gene expression (*n =* 3-4 differentiations or independent human islet donors/group). (J) Minimum to maximum box and whisker plots of motif deviation z-scored across SC-β cells differentiation trajectory and primary β cells generated from publicly-available scATAC-seq data^28^ representing PPARα motif activity (*n =* 3-4 differentiations or independent human islet donors/group).

We next wished to determine the mechanisms by which PPAR transcriptional pathways are lower in SC-islets. PPAR⍺ classically complexes with RXR, and together with PPAR and RXR ligands and transcriptional co-activators, form a transcriptional complex that can bind PPAR response elements to facilitate target gene expression^49^. Thus, we first assessed expression of the PPAR and RXR transcriptional machinery in stage 6 SC-islets. We found that *PPARA*, *PPARG*, *RXRA*, and *RXRB* expression levels were higher in stage 6 SC-islets compared to human islets (Fig. 6D). While we found that PPARα protein levels appeared to rise throughout differentiation, PPARα protein levels appeared unchanged in stage 6 SC-islets compared to human islets (Fig. 6E). The expression of the PPAR-related co-activators *NCOA1*, *EP300*, *EIF4E*, *SRC*, and *CTNND1* were also more highly expressed in stage 6 SC-islets compared to the human islets (Fig. 6D). To determine if these findings also occurred specifically in β cells, we reanalyzed a publicly-available single-cell RNA-seq data set^28^ and found that expression of the *PPARA* and *RXRA* were significantly higher in SC-β cells compared to primary human β cells (Fig. 6F). Further, while we observed lower expression of (*PPARG1A*/*PGC1⍺*) in SC-islets, we did not observe differences in *PGC1*⍺ expression between SC-β cells and primary human β cells (Fig. 6D and F). To assess if chromatin accessibility at putative PPAR- and RXR-binding sites was affected in SC-β cells, we next reanalyzed publicly-available single-cell ATAC-seq data from SC-β cells and primary human β cells^28^. We found that PPAR and RXR motifs were highly accessible in SC-β cells compared to primary β cells (Fig. 6G). These results suggest that limitations in the PPAR⍺/γ transcriptional program in SC-β cells was not due to a loss of the PPAR transcriptional machinery or chromatin accessibility.

We next identified a differentiation trajectory of stage 6 SC-β cells using publicly-available single-cell RNA-seq and single-cell ATAC-seq data^28^ to understand the dynamics of PPAR⍺ expression and activity during β cell differentiation. Using single-cell RNA-seq expression data specifically from stage 6 SC-islet cells, we first identified a developmental trajectory from immature *INS*^+^ β cells co-expressing the key endocrine progenitor transcription factor *FEV* to more mature β cells with higher *INS* expression and low/absent expression of *FEV* (Fig. S3A-B). Stage 6 SC-β cell trajectory was then extracted and combined with single-cell RNA-seq data from primary human β cells to compare expression. We then evaluated several β cell transcriptional regulators, observing greater expression in *PDX1* and *PAX6* across the SC-β cell differentiation trajectory (Fig. S4A). Similarly, *PPAR*⍺ expression increased along the SC-β cell differentiation trajectory (Fig. 6H). Despite this, we observed that the PPAR⍺ gene signature score, calculated using the expression of 51 known PPAR⍺ transcriptional targets^50^, remained at a lower level than primary human β cells throughout the SC-β cell differentiation trajectory (Fig. 6I). We next used ChromVAR^51^, a computational approach to estimate a per-cell bias-corrected measure of the change in chromatin accessibility in sets of peaks containing the same transcription factor motif compared to an average cell profile, which serves as a proxy for motif activity. Again, an evaluation of known β cell transcriptional activators revealed increases in *NEUROD1* and *PAX6* motif deviation across SC-β cell differentiation trajectory (Fig. S4B). Intriguingly, we observed that the estimated PPAR⍺ motif deviation did not change significantly across the SC-β cell differentiation trajectory but was consistently greater than that of primary β cells (Fig. 6J). Taken together, our analysis of bulk RNA-seq, pseudo-bulk single-cell RNA-seq and single-cell ATAC-seq, and transcriptomic and chromatin accessibility profiling across the SC-β cell differentitation trajectory, all revealed that SC-β cells possess lower expression of target genes within the PPAR⍺ pathway that do not appear to be due to reduced expression of the PPAR transcriptional machinery, accessibility of PPAR-binding motifs, or PPAR⍺ motif activity.

### WY14643 improves the differentiation of SC-β cells

While the actions of PPAR⍺/γ are well known in many cell types, including studies in postnatal β cells^52,53^, the importance of PPAR⍺/γ during β cell differentiation are unclear. We speculated that the limited activation of PPAR⍺/γ-related mitochondrial programming could be related to the lack of PPAR⍺/γ ligands during SC-islet differentiation. Notably, the primary ligand for RXR is retinoic acid, which is routinely included in SC-islet differentiation protocols^4,54^. Ligand binding to PPARs induces a conformational change that displaces co-repressors and enhances binding to co-activators to promote the expression of target genes^55,56^. PPAR agonists have also been shown to not be required to increase the number of PPARγ-binding sites, suggesting that chromatin accessibility at PPAR motifs in SC-β cells could be maintained and poised for activation in the absence of an agonist^57^, consistent with our observations above (Fig. 6F and I). Therefore, we hypothesized that addition of a PPAR agonist may improve SC-β cell differentiation.

As a proof of concept, we chose to include the potent PPAR⍺ agonist WY14643^58–61^ during stage 6 of the differentiation program given our observations of a relative lack of PPAR⍺ target gene expression throughout the SC-β cell maturation trajectory despite a maintenance of PPAR⍺ motif activity (Fig. 6I-J and 7A). Intriguingly, we found that WY14643 treatment significantly increased the percentage of C-peptide-positive SC-islets without changes to the glucagon-, somatostatin-, or polyhormone-positive cell populations (Figs. 7B). WY14643 treatment also significantly raised the frequency of NKX6.1 and C-peptide co-positive SC-β cells with similar increases in PDX1 and C-peptide co-positivity (Fig. 7C-D).

**Figure 7.**
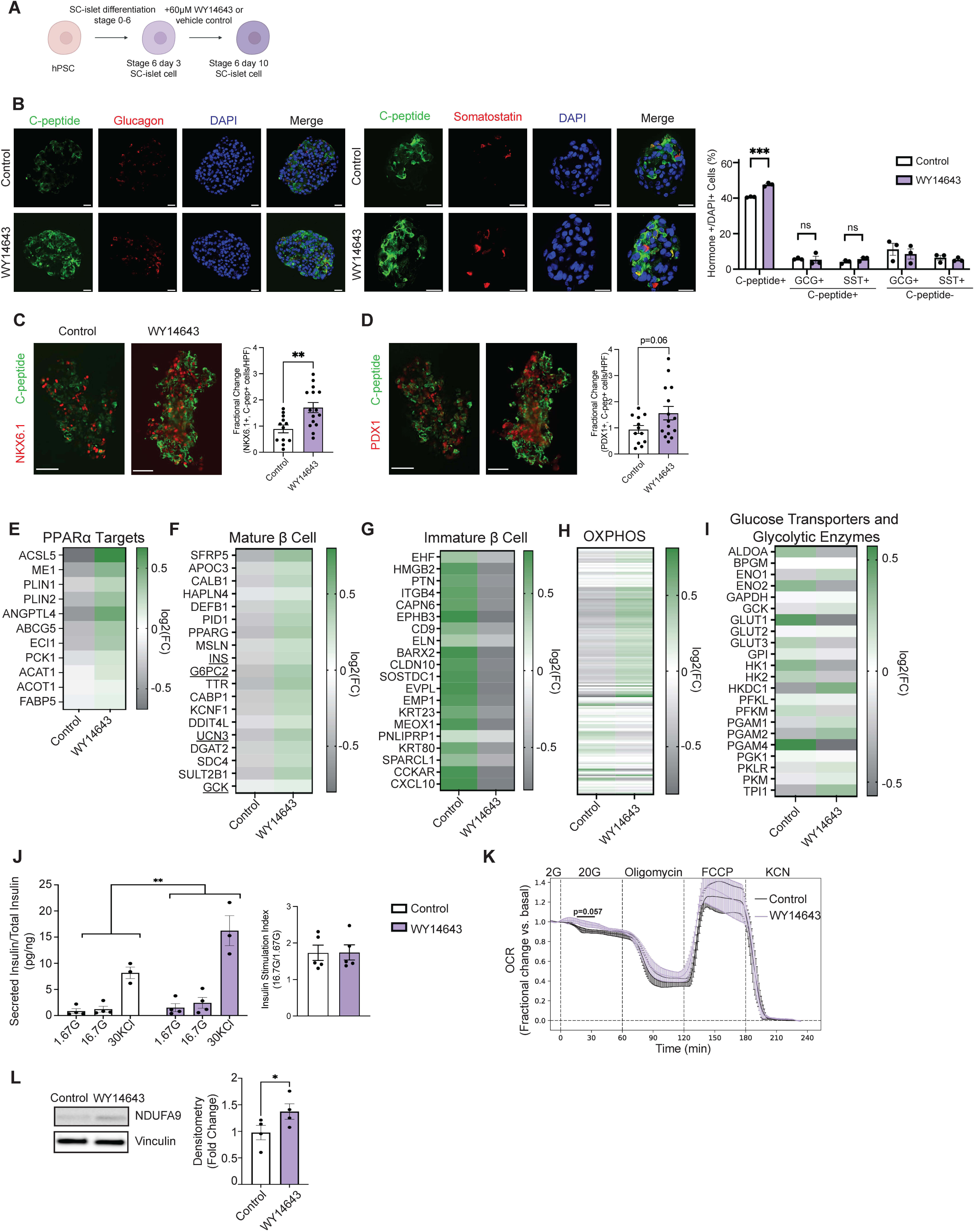
WY14643 promotes the formation of SC-β cells. (A) A schematic diagram illustrating use of the PPAR⍺ agonist WY14643 within the differentiation protocol. (B) Immunofluorescence images at 60X magnification (left) of vehicle control and WY14643-treated stage 6 SC-islets for C-peptide (green), glucagon (red), and DAPI (DNA - blue) or C-peptide (green), somatostatin (red), and DAPI (DNA - blue) (*n =* 3/group). Scale bars, 20 μm. Quantification (right) of C-peptide positive cells, dual hormone-positive cells, either C-peptide and glucagon (GCG) dual-positive or C-peptide and somatostatin (SST) dual-positive, as well as GCG-positive cells, and SST-positive cells (*n =* 3 differentiations/group, ∼1000 cells quantified/differentiation and respective condition); \*\*\**P <* 0.001 by unpaired Student’s *t*-test. (C) Immunofluorescence images at 20X magnification (left) of vehicle control and WY14643-treated stage 6 SC-islets for C-peptide (green) and NKX6.1 (red) (*n =* 3/group). Scale bars, 50 μm. Quantification (right) of the fractional change of NKX6.1/C-peptide co-positive cells per high powered field (*n =* 12-15 fields quantified/group, ∼1000 cells quantified/differentiation and respective condition from 3 differentiations/group); \*\**P <* 0.01 by unpaired Student’s *t*-test. (D) Immunofluorescence images at 20X magnification (left) of vehicle control and WY14643-treated stage 6 SC-islets for C-peptide (green) and PDX1 (red) (*n =* 3/group). Scale bars, 50 μm. Quantification (right) of the fractional change of PDX1/C-peptide co-positive cells per high powered field (*n =* 12-15 fields quantified/group, ∼1000 cells quantified/differentiation and respective condition from 3 differentiations/group). (E) Differential RNA expression heatmap (presented as integrated mean/group) of PPAR⍺ targets for vehicle control and WY14643-treated stage 6 SC-islets (*n =* 4/group). (F) Differential RNA expression heatmap (presented as integrated mean/group) of mature β cell markers for vehicle control and WY14643-treated stage 6 SC-islets (*n =* 4/group). (G) Differential RNA expression heatmap (presented as integrated mean/group) of immature β cell markers for vehicle control and WY14643-treated stage 6 SC-islets (*n =* 4/group). (H) Differential RNA expression heatmap (presented as integrated mean/group) of genes encoding OXPHOS components for vehicle control and WY14643-treated stage 6 SC-islets (*n =* 4/group). (I) Differential RNA expression heatmap (presented as integrated mean/group) of genes encoding glucose transporters and glycolytic enzymes for vehicle control and WY14643-treated stage 6 SC-islets (*n =* 4/group). (J) Insulin secretion (left) with insulin stimulation index (right) following static incubations in 1.67 mM glucose (1.67G), 16.7 mM glucose (16.7G) and 30 mM KCl (30KCl) normalized to insulin content in vehicle control and WY14643-treated stage 6 SC-islets (*n =* 4 differentiations/group); \*\**P <* 0.01 by two-way ANOVA. (K) Oxygen consumption rate (OCR) fractional change in vehicle control and WY14643-treated stage 6 SC-islets (*n =* 3 differentiations) following exposure to 2 mM glucose (2G), 20 mM glucose (20G), 10 μM oligomycin, 1 μM FCCP, and 3 mM KCN. Data are presented as mean from independent differentiations ± SEM; *P* = 0.057 by ANOVA of fractional OCR value at designated time points following 20 mM glucose exposure. (L) Representative Western blot analysis of NDUFA9 protein expression for vehicle control and WY14643-treated stage 6 SC-islets (left). Vinculin serves as a loading control. Quantification (right) of NDUFA9 expression (normalized to Vinculin expression) via densitometry of the Western blots (*n =* 4/group); \**P <* 0.05 by unpaired Student’s *t*-test.

To gain further insight into the mechanisms underlying the expansion of SC-β cells upon PPAR agonism, we performed bulk RNA-seq studies on WY14643-treated SC-islets compared to control-treated SC-islets. Firstly, we observed the expected induction of a subset of PPAR⍺ targets in WY14643-treated SC-islets compared to control-treated SC-islets (Fig. 7E). We next observed higher expression of genes reported to be expressed in mature β cells^62^ upon WY14643 exposure, including *INS*, *UCN3*, *GCK*, and *G6PC2*, as well as lower expression of genes enriched in immature β cells^62^, which could also be reflective of a higher percentage of β cells overall in bulk assessments of WY14643-treated SC-islets (Fig. 7F-G). In addition, we found greater expression of ETC-OXPHOS-related genes, with minimal changes in expression of genes related to glucose uptake and glycolysis, in WY14643-treated SC-islets compared to control-treated SC-islets (Fig. 7H-I). Further, we found that addition of WY14643 significantly improved insulin secretion in SC-islets compared to controls, yet the fold induction of insulin secretion by glucose was similar between the groups (Fig. 7J). Together, these studies suggested that WY14643 improves the formation of SC-β cells.

We next evaluated potential mitochondrial mechanisms promoting SC-β cell formation following WY14643 exposure. We first found modest increases in mitochondrial respiration following high glucose stimulation in WY14643-treated SC-islets (Fig. 7K). We observed that WY14643 treatment resulted in greater protein expression of the complex I subunit NDUFA9 compared to controls, however, changes in other OXPHOS subunits were not found (Fig. 7L and S5A). We also observed no changes in mitochondrial mass, expression of regulators of mitochondrial dynamics, or mitochondrial ultrastructure following WY14643 exposure (Figs. 7L and S5B-D). We next found that that WY14643 exposure reduced both fatty acid and acylcartinine concentrations in SC-islets, consistent with the induction of fatty acid oxidation following PPAR⍺ agonism (Fig. S6A-B). To evaluate the contributions of mitochondrial pyruvate transport or long chain fatty acid oxidation to the effects of WY14643 on SC-β cell formation, we exposed stage 6 SC-islets to WY14643 (or vehicle control) for 7 days in the presence or absence of the mitochondrial pyruvate carrier (MPC) inhibitor UK5099 or the CPT1A inhibitor etomoxir. Indeed, UK5099 exposure abrogated the effects of WY14643 to increase the frequency of SC-β cells, while etomoxir exposure increased the presence of GCG+ SC-⍺ cells (Fig S6E-F). Finally, we examined other metabolic signaling pathways, including Calcium/calmodulin-dependent protein kinase II (CaMKII), AMP-activated protein kinase (AMPK), and acetyl co-A carboxylase (ACC), yet did not observe differences in the phosphorylation and activation of these pathways following WY14643 exposure (Figure S6G-I). Taken together, these results suggest that WY14643 induces both mitochondrial oxidative and fatty acid metabolism to promote SC-β cell formation.

We next wanted to determine if the beneficial effects of WY14643 were visible in other SC lines and also durable *in vivo* following transplantation. We first tested the effects of WY14643 in stage 6 SC-islets differentiated from the 19-9-11 induced pluripotent stem cell (iPSC) line (Figure S7A-B). We observed that addition of WY14643 for 7 days increased the insulin secretion index of iPSC-derived SC-islets (Fig. S7C). Next, we treated SC-islets (from the H1 ESC line) with WY14643 or vehicle control for 7 days *in vitro* prior to seeding SC-islets on a poly(lactide-*co*-glycolide) (PLG) microporous scaffold prior to transplantation into the epididymal fat pad of immunodeficient NOD-*scid* IL2Rg^null^ (NSG) mice. We delivered SC-islet grafts to the epididymal fat pad rather than the renal subcapsular space as the epididymal fat pad is a translationally relevant transplant site^63,64^ (Fig. 8A). We then retrieved the scaffolds 10 days posttransplant due to the limited half-life of WY14643, and we did not administer WY14643 *in vivo* to avoid systemic effects of the agonist. Following graft retrieval, we observed SC-islets appeared as expected within the scaffold with recipient cell growth within and surrounding the xenograft (Fig. 8B). Notably, we observed a durable effect of WY14643 on the SC-β cell lineage 10 days post-transplantation, with a significant increase in C-peptide-positive SC-β cells compared with vehicle-treated controls (Fig. 8C-D). Further, we observed that WY14643 did not affect the percentage of somatostatin- or glucagon-positive cells but did lead to a modest reduction in the immature polyhormone-positive cell population post-transplantation (Fig. 8C-D). Moreover, WY14643 treatment led to a significant increase in insulin content within the explanted scaffold graft compared to vehicle-treated controls (Fig. 8E). We also observed that WY14643 treatment did not significantly alter the frequency of Ki-67/C-peptide co-positive cells, suggesting WY14643 likely did not enhance SC-β cell replication (Fig. 8F). WY14643 treatment also did not alter mitochondrial morphology in SC-β cells post-transplantation (Fig. 8G). Taken together, the addition of a PPAR⍺ agonist to stage 6 SC-islets improved the formation of SC-β cells, which was durable *in vivo* post-transplantation.

**Figure 8.**
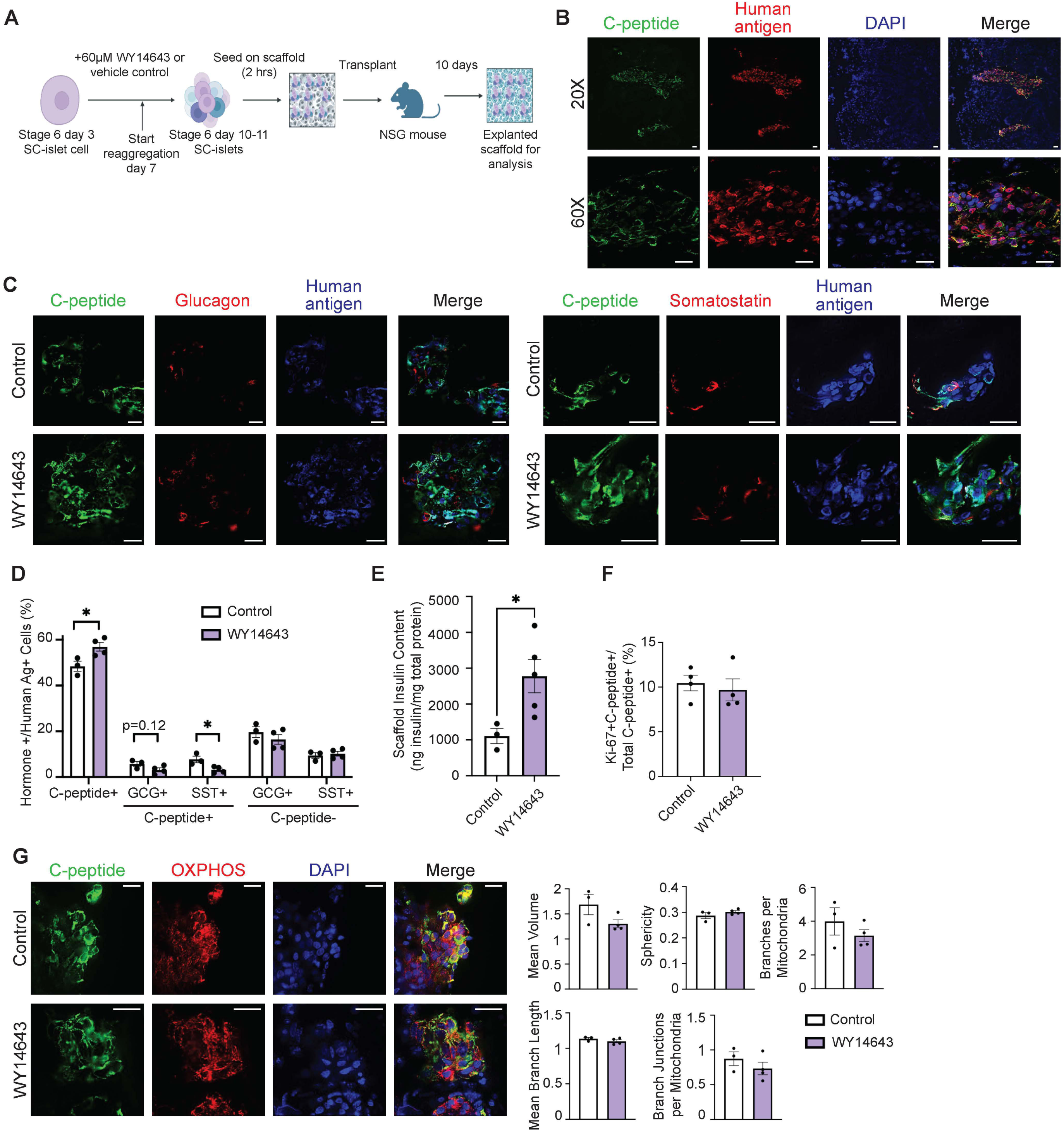
Formation of SC-β cells by WY14643 is durable *in vivo* following transplantation. (A) A schematic diagram illustrating the addition of WY14643 to the differentiation protocol, seeding of SC-islets onto poly(lactide-*co*-glycolide) (PLG) microporous scaffolds, and scaffold transplantation into NSG mice. (B) Immunofluorescence images at 20X and 60X magnification of stage 6 SC-islets for C-peptide (green), human nuclear antigen (red), and DAPI (blue) (*n =* 3/group). Human nuclear antigen was used to distinguish human transplanted cells from murine recipient cells in the inguinal fat pad. Scale bars, 20 μm. (C) Immunofluorescence images at 60X magnification of vehicle control and WY14643-treated stage 6 SC-islets in explanted scaffold from NSG mice 10 days post-transplant for C-peptide (green), glucagon (red), and DAPI (blue) or C-peptide (green), somatostatin (red), and DAPI (blue) (*n =* 3-4/group). Scale bars = 20 μm. (D) Quantification of C-peptide-positive cells only, dual hormone-positive cells, both C-peptide- and GCG- or SST-positive cells, GCG-positive cells, and SST-positive cells by immunostaining in vehicle control and WY14643-treated stage 6 SC-islets in explanted scaffold from NSG mice 10 days post-transplant (*n =* 3-4/group); \**P <* 0.05 by unpaired Student’s *t*-test. (E) Insulin content normalized to total protein content for vehicle control and WY14643-treated explanted scaffolds from NSG mice 10 days post-transplant (*n =* 3-4/group); \**P <* 0.05 by unpaired Student’s *t*-test. (F) Quantification of Ki-67 and C-peptide co-positive cells by immunostaining in vehicle control and WY14643-treated stage 6 SC-islets in explanted scaffold from NSG mice 10 days post-transplant (*n =* 4/group). (G) Immunofluorescence images (right) at 60X magnification of vehicle control and WY14643-treated stage 6 SC-islets in explanted scaffold from NSG mice 10 days post-transplant for C-peptide (green), OXPHOS (red), and DAPI (blue) (*n =* 3-4/group). Scale bars, 20 μm. Quantification (right) of mitochondrial morphology and networking utilizing Mitochondria Analyzer (*n =* 3-4/group, ∼150 cells analyzed per differentiation and treatment).

## DISCUSSION

Here, we reveal a vital importance for mitochondrial programming to promote the differentiation of SC-β cells. Our studies elucidate specific mechanisms underlying limitations in SC-β cell mitochondrial metabolism, including reductions in expression and function of the ETC-OXPHOS system and mitochondrial fatty acid metabolism. We observe limitations in glucose-stimulated respiration in SC-islets without alterations in mitochondrial mass, structure, mtDNA content, and mtDNA replication. We identify several candidates that may contribute to limitations in mitochondrial programming during SC-β cell maturation, notably the nuclear receptors PPAR⍺ and PPARγ. We demonstrate that the PPAR⍺ and PPARγ transcriptional machinery is intact and chromatin at the sites of PPAR⍺ and PPARγ binding motifs is accessible, but expression of PPAR⍺ and PPARγ target genes is reduced, suggesting a deficiency of PPAR ligands in the SC-islet differentiation process. To test a proof of concept of the importance of PPAR-mediated programming in SC-β cells, we found that the PPAR⍺ agonist WY14643 improved mitochondrial respiration, fatty acid oxidation, and importantly, insulin secretion. Moreover, WY14643 increased the formation of C-peptide-positive SC-β cells both *in vitro* and upon transplantation within a translationally relevant engraftment site, as well as enhancing insulin content within SC-islet grafts. Thus, our data reveal the potential for targeting PPAR⍺-dependent mitochondrial programming to improve SC-β cell differentiation, expanding beyond classical roles for mitochondria simply as a client organelle necessary for β cell bioenergetics.

Our results suggest that the activation of mitochondrial transcriptional programs can expand the SC-β cell lineage. While the synchronistic connections between mitochondria and cell fate are well appreciated in other tissue types, such as brown adipocytes^65,66^, this connection has not been clear during β cell differentiation. Interestingly, PPAR⍺/γ regulate the differentiation of adipocytes, keratinocytes, bone, and several immune cell types and promote the expression of genes related to lipid metabolism and mitochondrial biogenesis^67–73^, yet their function in SC-β cell differentiation had not previously been observed. In adult human islets, PPAR⍺ overexpression or treatment with a PPAR⍺ agonist offered protection from lipotoxicity and preserved GSIS^74,75^. Similarly, PPARγ agonists preserved β cell function in humans with T2D and protected human islets from lipotoxicity^52,76–78^. However, a role for mitochondrial programming by PPAR⍺/γ to promote human β cell differentiation was not examined. Our studies reveal the potential for targeting PPAR⍺ to improve mitochondrial respiration and fatty acid metabolism and expand the SC-β cell lineage. Indeed, we speculate that PPAR⍺ could act through mitochondria to promote β cell differentiation by supporting the bioenergetic demands of differentiation or through increased substrate availability to facilitate histone modifications and chromatin accessibility at key β cell differentiation genes, which remains to be tested in the future. Future work will also be needed explore the addition of PPAR agonists in other SC-β cell differentiation stages, define the roles of PPARs on metabolism throughout β cell development, and to explore the potential for shared PPAR-related mechanisms underlying differentiation across tissues, including SC-β cells as well as adipocytes, keratinocytes, bone, and immune cells.

PPAR⍺/γ are well known regulators of lipid metabolism, and PPAR⍺ is a central regulator of fatty acid oxidation. Our studies suggest that WY14643 enhances both mitochondrial fatty acid oxidation and respiration in SC-islets. Beyond their roles in lipid metabolism and mitochondrial biogenesis, emerging studies demonstrate crosstalk between ERR⍺ and PPAR⍺, which could suggest the potential that WY14643 might indirectly support expression of OXPHOS targets through ERRs^44,79^. Our studies define a previously unknown impairment in mitochondrial lipid metabolism in SC-islets with beneficial roles of WY14643 on SC-β cell formation, as well as OXPHOS gene expression, which appears distinct from the roles of ERRγ to improve SC-β cell GSIS through induction of OXPHOS and suppression of glycolysis^11^. Future studies will be crucial to discriminate both the specific and integrated contribution of each of these highly integrated mitochondrial transcriptional regulators on SC-islet development, including PPAR⍺, PPARγ, ERRγ, and PGC1⍺. The impact of pharmacologic agonists of ERRγ^80^ and specific, partial, or dual agonists of PPAR⍺ and PPARγ^81–83^, to together improve both SC-β cell differentiation (through PPAR⍺) and maturation/GSIS (through ERRγ) will be intriguing to assess for future translational benefit.

Our studies expand upon the metabolic differences observed in SC-islets compared to primary human islets, which have understandably focused to date on glucose metabolism given the pivotal importance of glucose metabolism for β cell insulin secretion. We found that SC-islets do not increase mitochondrial respiration upon high glucose stimulation, similar to the observations of others^6,10^. We noted that SC-islets exhibited both lower oxygen consumption following oligomycin exposure and a higher peak oxygen consumption after the uncoupler FCCP compared to human islets, which had not been observed previously. These effects could be a function of differences of tissue penetration of pharmacologic agents between SC-islets and human islets despite their similar size and appearance. Alternatively, the differences in SC-islet responses to oligomycin and FCCP could be indicative that SC-islets may be more coupled or possibly have reductions in proton leak compared to primary human islets. Primary β cells are known to be remarkably uncoupled, and blockade of adenosine nucleotide translocase (ANT)-mediated basal proton leak impairs GSIS^84–87^. Indeed, SC-islets express significantly lower levels of ANT than primary human islets (log_2_FC −0.66, FDR < 0.05). We also observe alterations in mitochondrial acylcarnitine profiles in SC-islets, indicative of limitations in mitochondrial lipid metabolism that may originate due to limitations in mitochondrial programming by PPAR⍺/γ. A recent study demonstrated that elevations in the sphingolipid ceramide limited the maturation of SC-β cells, suggesting that other lipid subtypes may play distinct roles in differentiation^88^. Reduced levels of C3 and C5 acylcarnitines may also reflect alterations in branched chain amino acid metabolism^89^, an effect we also observed in SC-islets (Fig. 3D). Notably, fluctuations in amino acid availability at birth may also impact the nutrient sensitivity of mTORC1 signaling pathway to affect β cell maturation^90^. Unsaturated fatty acids can serve as endogenous ligands for PPAR⍺ and/or PPARγ, leading us to speculate a possible similar role for these metabolites in β cell maturation, especially given the fatty acid composition of maternal breast milk^91–94^. It will be intriguing to test if future approaches to modulate β cell uncoupling or nutrient exposure will have a beneficial impact on SC-β cell differentiation and maturation. On the other hand, exploration of the SC-islet microenvironment or culture conditions, could provide an alternative approach to clarify the metabolic control of SC-β cell differentiation and function.

Limitations in our study included use of RNA-seq data performed on bulk SC-islet differentiations, which have a different cell type composition than bulk primary human islets^28^. Our conclusions were supported, however, by re-analysis and inclusion of published single cell RNA-seq of SC-β cells and primary β cells^28^. Further, despite reports of mitochondrial dysfunction in SC-islets from numerous groups, the effect of PPAR agonists will need to be evaluated in other SC-β cell differentiation protocols, as our observations could be protocol-dependent. While numerous studies, including genetic knockout models, indicate a specific role for WY14643 both *in vitro* and *in vivo* through PPAR⍺^59–61^, WY14643 has been reported to have weak PPARγ activity *in vitro*^95^. Thus, additional studies to dissect the contribution of PPARγ on SC-islet differentiation would be valuable in the future. An additional limitation of our study is the need to determine if PPAR agonism, such as by WY14643 or other agents, is able to promote SC-β cell differentiation and function to ultimately improve glycemic control upon engraftment in diabetic mice *in vivo*. We also did not explore the role for systemic administration of PPAR agonists post-transplantation, which might provide additional opportunities to enhance *in vivo* SC-β cell function.

Our findings support metabolic limitations previously found in the generation of SC-β cells, but also open new roles for of PPAR⍺ and mitochondria in the regulation of SC-β cell differentiation. Our results reinforce the findings of several studies of SC-β cell metabolism, including defects in OXPHOS expression, persistence of aberrant glycolytic metabolism, and limitations in expression of the ERR transcriptional pathway^6,10,11^, while clarifying new roles for mitochondrial fatty acid metabolism and OXPHOS mediated at least in part by PPAR⍺. Our results also demonstrate the novel potential to activate PPAR⍺ as a highly druggable target to improve SC-β cell differentiation. Taken together, our results corroborate limitations that remain in the formation of SC-β cells and identify new potential routes to overcome these deficits, possibly requiring pharmacologic targeting of several complementary pathways to generate higher quantities of fully functional SC-β cells for enhanced therapeutic use in diabetes. Early clinical trials have provided excitement regarding the therapeutic potential for SC-islet transplantation to provide a functional cure for T1D^3,96^. However, these allografts require up to 6 months to optimize their function, and the long-term viability of these cells remain unclear^3^. Thus, our studies highlight the opportunity to focus on mitochondrial programming pathways to improve the differentiation, metabolism, and function of SC-β cells to more closely resemble primary β cells as an approach to improve durable cell-based therapies for T1D.

## EXPERIMENTAL MODEL AND SUBJECT DETAILS

### Stem cell-derived islets (SC-islets)

SC-islets were differentiated from H1, HUES8, and IPS-DF 19-9-11 T.H (feeder independent; WiCell) human pluripotent stem cells (hPSCs) using a 6-stage protocol, as reported previously^4,63,64,97^. Briefly, undifferentiated cells were dispersed into single cells, and 5 × 10^6^ cells were seeded in 6-well cell culture treated plates coated with Matrigel (Corning). Cells were cultured for 24 h in mTeSR1 and then cultured in the differentiation media for 6 stages. During each stage, cells received daily media changes with stage-specific basal media supplemented with growth factors and small molecules to induce differentiation. The stages of this protocol are defined as definitive endoderm (stage 1), primitive gut tube (stage 2), pancreatic progenitor (stages 3 and 4), endocrine progenitor (stage 5) and immature stem cell derived β-cell (stage 6). At the end of each stage, cells could be dissociated from the plate, fixed and analyzed using intracellular flow cytometry for stage-specific biomarkers. For reaggregation from planar culture, cells were dispersed into single cells using TrypLE and seeded onto suspension culture 6-well plates at 8 × 10^6^ cells per well. Y27632 was included in the media for the first 24 hours of culture, then media was changed every other day for the duration of culture. Plates were placed on an orbital shaker set to 100 rpm inside a humidified incubator. Reaggregted cell clusters were treated with 60 μM WY14643 alone or with 4 μM UK5099 or 25 μM Etomoxir days 3-10. For scaffolds, reaggregated cell clusters were collected and seeded on the scaffold in the same manner as described above after 3–4 days in culture,.

### Microporous scaffold fabrication

Poly(lactide-*co*-glycolide) (PLG) microporous scaffolds were fabricated as previously described^63^. Briefly, PLG microporous scaffolds were fabricated by compression molding PLG microspheres (75 : 25 mole ratio d,l-lactide to glycolide) and 250 to 425 μm salt crystals in a 1:30 ratio of PLG microspheres to salt. The mixture was humidified in an incubator for 7 minutes. Scaffolds were compression molded with 77.5 mg of polymer–salt mixture into cylinders 5 mm in diameter by 2 mm in height using a 5 mm KBr die (International Crystal Laboratories, Garfield, NJ) at 1500 psi for 30 seconds. Molded constructs were gas foamed in 800 psi carbon dioxide for 16 hours in a pressure vessel. The vessel was depressurized at a controlled rate for 30 min. On the day of cell seeding, scaffolds were leached in water for 1.5 hours, changing the water once after 1 hour. Scaffolds were disinfected by submersion in 70% ethanol for 30 seconds and rinsed multiple times with phosphate-buffered saline (PBS).

### Scaffold seeding of SC-islets

To initiate scaffold culture differentiations from reaggregated clusters, reaggregated clusters were seeded on scaffolds at a density of 125 × 10^6^ cells/cm^3^. Prior to seeding, scaffolds were washed in cell media solution then briefly dried on sterile gauze to improve the absorption of the cell solution into the scaffold. Cells were distributed across both faces of the scaffold and then incubated for 2 hours to allow cell solution to be further absorbed into the scaffold before differentiation media was added.

### Human islets

All human samples were procured from de-identified donors without diabetes from the Integrated Islet Distribution Program, Alberta IsletCore, or Prodo Laboratories and approved by the University of Michigan Institutional Review Board. Human primary islets were cultured at 37°C with 5% CO_2_ in PIM(S) media (Prodo Laboratories, Aliso Viejo, CA, USA) supplemented with 10% FBS, 100 U/mL penicillin/streptomycin, 100 U/mL antibiotic/antimycotic, and 1 mM PIM(G) (Prodo Laboratories, Aliso Viejo, CA, USA). Islets were used from male and female donors, and donor information is provided in Table S1.

### Transplantation of reaggregated SC-islets in immunodeficient mice

SC-islets were transplanted as previously described^63^. All mice were maintained in accordance with the University of Michigan’s Institutional Animal Care and Use Committee under specific pathogen-free conditions and were assigned randomly to different experimental groups. Up to 5 mice were housed per cage and were maintained on regular chow with *ad libitum* access to food and water on a 12 h light-dark cycle. 8–10-week-old male NOD.Cg-Prkdc^scid^ Il2rgtm1Wjl/SzJ (NSG) mice (Jackson Laboratories) were used for transplantation studies. On the day of transplantation, the abdomen was shaved and sterilized. A small incision was made in the peritoneal wall and the epididymal fat pads were unwrapped outside the body. As described above, reaggregated cell clusters were collected and seeded on both faces of the scaffold and incubated at least 2 h before transplantation. Reaggregated cell-laden scaffolds were placed on the fat pads, wrapped in fat pad tissue, and placed back into the peritoneal cavity. Each mouse received approximately 5 million cells in clusters on each scaffold, equaling a total of 10 million cells transplanted per mouse. Mice were sutured, surgically stapled, and received carprofen (0.5 mg/mL) the day of and after surgery. Body weight was measured daily for 10 days prior to euthanasia and subsequent graft harvest.

## METHOD DETAILS

### Culture methods

Human pluripotent stem cell differentiation in H1 and HUES8 cells was performed as per the protocol previously published by Hogrebe, et al. 19-9-11 T.H (feeder independent) induced pluripotent cells were differentiated through stage 4 using the STEMdiff™ Pancreatic Progenitor Kit (#05120) and subsequently as above.

### Flow cytometry

Single cell dissociations were fixed in 4% paraformaldehyde and blocked with 5% normal donkey serum in 0.1% Triton-X PBS. Cells were incubated overnight with primary antibodies, washed and subsequently incubated with appropriate secondary antibodies. Stained, fixed cells were analyzed by ZE5 cell analyzer (Bio-Rad). Primary and secondary antibodies used: anti-OCT-3/4 (Santa Cruz Biotechnology, catalog # sc-5279), anti-chromogranin A (Invitrogen, catalog # PIMA533052), anti-C-peptide (Developmental Studies Hybridoma Bank, University of Iowa, catalog # GN-ID4-S), anti-NKX6–1 (Developmental Studies Hybridoma Bank, University of Iowa, catalog # F55A12-S), anti-PDX1 (R&D Systems, catalog #AF2419), AlexaFluor 488 (Invitrogen, catalog # A21208), AlexaFluor 488 (Invitrogen, catalog # A21202), AlexaFluor 647 (Invitrogen, catalog # A31571), AlexaFluor 647 (Invitrogen, catalog # A21447), AlexaFluor 647 (Invitrogen, catalog # A31573).

### Static *in vitro* insulin secretion measurements

Static insulin secretion assays on human islets and SC-islets were performed following 1 h incubations with 1.67 mM and 16.7 mM glucose, and 30mM KCl in HEPES-supplemented KRB buffer containing 135 mM NaCl, 4.7 mM KCl, 1.2 mM KH_2_PO_4_, 5 mM NaHCO_3_, 1.2 mM MgSO_4_.7H_2_O, 1 mM CaCl_2_, 10 mM HEPES and 0.1% BSA (pH 7.4). Samples were analyzed for insulin release using Human Insulin Chemiluminescence ELISA (ALPCO; Salem, NH, USA) as per the manufacturer’s protocol.

### Insulin content of explanted scaffolds

Explanted scaffolds were placed in 9.1 mM HCl 95% EtOH solution. For extraction, samples were sonicated 5x for 15 sec, resting on ice in between sonication periods. Samples were rotated at 4°C overnight. Samples were then spun at 2500 RPM for 30 min at 4°C. Supernatants were stored at −80°C until analysis. Samples were analyzed for insulin release using Human Insulin Chemiluminescence ELISA (ALPCO; Salem, NH, USA) as per the manufacturer’s protocol.

### Mitochondrial respirometry

The oxygen consumption rate was measured in stage 6 SC-islets and primary human islets using the BaroFuse (EnTox Sciences, INC; Mercer Island, WA, USA) as previously described^20^. 200 islets were placed in Krebs-Ringer Bicarbonate HEPES (KRBH) buffer with a 2 mM glucose per condition. Islets were subject to the following conditions: 20 mM glucose, 10 μM oligomycin (Sigma), 1 μM FCCP (Sigma-Aldrich), and 3 mM KCN (Thermo Scientific) for 60 min each.

### Metabolomics

Cells were cultured according to Hogrebe et al^4^. On the day of sample collection, samples were incubated in MCDB 131 with 10.5 g BSA, 5.2 mL GlutaMAX, 5.2 mL P/S, 5 mg heparin, 5.2 mL MEM nonessential amino acids (Corning, 20–025-CI), 84 μg ZnSO4 (MilliporeSigma, 10883), 523 μL Trace Elements A (Corning, 25–021-CI), and 523 μL Trace Elements B (Corning, 25–022-CI) and either 5.5mM or 20mM glucose for 3 hr. Media was removed, and the cells were washed 1X with ice-cold 150 mM ammonium acetate in LC-MS grade water. Cells were harvested with 200 μL of ice-cold methanol and frozen at −80°C until sample preparation. For extraction, 200 μL of cold water was added to the cells and the cells were scraped. The resulting homogenate was sonicated on ice for 10 sec. 400 μL of chloroform was added. Samples were centrifuged at 17,000 x *g* for 10 min and the resulting top layer was collected and taken to dryness under nitrogen. Samples were reconstituted in 30 μL of 2:1 ACN: water, filtered and 5 μL were injected for analysis. The samples were separated as previously described^98^. LC-MS analysis was performed on the Agilent 6546 quadrupole TOF mass spectrometer, which was coupled to Agilent 1290 LC. The gas temperature was set at 225°C, the drying gas flow at 10 L/min, the nebulizer at 40 psi, the sheath gas temperature at 300°C, and the sheath gas flow at 12 L/min. The Fragmentor (Agilent) was set at 125 V, the skimmer at 65 V, and VCap at 3,000 V. For verification of metabolite identity and retention time, authentic standards of all measured metabolites were run separately and spiked into pooled samples. Metabolites were separated on an InfinityLab Poroshell120 HILIC-Z, 2.7 μm, 2.1 × 150 mm column (Agilent Technologies, CA, USA). Mobile phase A was composed of 90% 10 mM ammonium acetate pH 9.0 with ammonia, 10% acetonitrile (ACN) with 2.5 uM InfinityLab Deactivator Additive (Agilent Technologies, CA, USA) and mobile phase B was composed of 15% 10 mM ammonium acetate pH 9.0 with ammonia, 85% acetonitrile with 2.5 uM InfinityLab Deactivator Additive. The flow rate was 0.25 mL/min, and the gradient was as follows: 0-2 min at 95 % B, 2-5 min at 95% B, 5-5.5 min at 86%, 5.5-8.5 min at 86% B, 8.5-9 min at 84% B, 9-14 min at 84% B, 14-17 min at 80% B, 17-23 min at 60% B, 23-26 min at 60% B, 26-27 min at 95% B, and 27-35 min at 95% B. Column compartment temperature was kept at 25 °C. Data were acquired in negative mode.

### Lipidomics

Media were removed, and metabolites were extracted with solvent (8:1:1 ratio of methanol/Chloroform/water) containing stable isotope-labeled internal standards. As previously described^99^, an Agilent 6410 triple quadrupole MS system equipped with an Agilent 1200 LC System and an ESI source was utilized. Metabolite separation was achieved using gradient elution on a reverse phase XBridge C18 Column (50 mm × 2.1 mm, 2.5 μm, Waters, Milford, MA, USA) with the corresponding guard column (5 mm x 2.1 mm, 1.7 μm) maintained at 40 °C. LC vials were maintained at 4 °C in a thermostatic autosampler, and the injection volume was set at 4 µL. The mobile phase consisted of solvent A, 5 mM Ammonium Acetate, and solvent B, Acetonitrile. The flow rate was 0.25 mL/min. The gradient elution program was as follows: 0–8.50 min, 2% B; 1.5–9 min, linear gradient from 2% to 50% B; 9–14 min, linear gradient from 50% to 95% B; hold at 95% B for 3 min. The flow rate was 300 μL/min. Acylcarnitine species were each detected by their characteristic LC retention time in the MRM mode following ESI and comparing relative areas with those of corresponding standards. Concentrations of carnitine, acetylcarnitine (C2), propionylcarnitine (C3), butyrylcarnitine (C4), isovalerylcarnitine (C5), hexanoylcarnitine (C6), octanoylcarnitine (C8), myristoylcarnitine (C14), palmitoylcarnitine (C16), and oleoylcarnitine (C18) were calculated by ratios of peak areas of known concentrations of stable isotopically-labeled analogs.

### mtDNA content and replication assessments

25 human islets per donor were handpicked and washed twice with 1X PBS. 100,000 cells from differentiation stage 0 through 6 were washed twice with 1X PBS. Human islets and stem cells were pelleted and stored at −80°C until DNA extraction using the Blood/Tissue DNeasy kit (Qiagen; Germantown, MD, USA) as per manufacturer’s instructions. Relative mtDNA content was quantified by qPCR using mtDNA specific primers ND1 (5’-ACCCCCTTCGACCTTGCCGA-3’ and 5’-GGGCCTGCGGCGTATTCGAT-3’), and nuclear DNA specific primers for GAPDH (5’-AATCCCATCACCATCTTCCA-3’ and 5’-TGGACTCCACGACGTACTCA-3’) as previously described^100^. First-strand mtDNA replication levels were measured by qPCR following digestion by Mnl I, which site specifically digest double-stranded DNA^101,102^. Briefly, primers specific to 7 S mtDNA (5′-ATCAACTGCAACTCCAAAGCCACC-3′ and 5′-TGATTTCACGGAGGATGGTGGTCA-3′), whose amplicon contains a Mnl I site, were used to quantify relative single-stranded mtDNA forming the D-loop. Single-stranded DNA generated from the extension of 7S region was measured using primers specific to CYTB (5′-AGTCCCACCCTCACACGATTCTTT-3′ and 5′-AGTAAGCCGAGGGCCTCTTTGATT-3′), which also contains an Mnl I site. Primers specific to COXI (5′-ACCCTAGACCAAACCTACGCCAAA-3′ and 5′-TAGGCCGAGAAAGTGTTGTGGGAA-3′), whose amplicon does not contain an Mnl I site, were used to quantify non-replicating double-stranded mtDNA. The ratio of 7S or CYTB amplicons to the COXI amplicon were then calculated after qPCR of Mnl I-digested DNA to determine the relative amount of single-stranded mtDNA at the D-loop strand or initiation of first strand replication, respectively.

### Citrate synthase activity assay

1,000 human islets per donor were handpicked and washed twice with 1X PBS. One million cells from differentiation stage 0 through 6 were washed twice with 1X PBS. Human islets and stem cells were lysed with radioimmunoprecipitation assay buffer containing protease and phosphatase inhibitors (Calbiochem), and insoluble material was removed by centrifugation. Lysates were used to determine citrate synthase activity with the MitoCheck Citrate Synthase Activity Assay Kit (Cayman Chemical, Catalog# 701040) per manufacturer’s instructions.

### Western blotting

1,000 human islets per donor were handpicked and washed twice with 1X PBS. One million cells from differentiation stage 0 through 6 were washed twice with 1X PBS. Human islets and stem cells were lysed with radioimmunoprecipitation assay buffer containing protease and phosphatase inhibitors (Calbiochem), and insoluble material was removed by centrifugation. Equal amounts of proteins were resolved on 4%–20% gradient Tris-glycine gels (Bio-Rad) and transferred to nitrocellulose membranes (Bio-Rad). Membranes were then blocked in 5% milk for 1 h and immunoblotting was performed using antibodies to Vinculin (1:1000; Millipore, Catalog# CP74), TOM20 (1:1000; Cell Signaling Technology, Catalog# 42406), Total OXPHOS Human Antibody Cocktail (1:1000; Abcam, ab110411), NDUFA9 (1:1000; Abcam; ab14713), UQCRC2 (1:1000; Abcam; ab14745), PPARα (1:3000; ProteinTech; 66826-1), phospho-CAMKII (1:1000; Cell Signaling; 12716), CAMKII (1:50; Santa Cruz; sc-5306), phospho-AMPK (1:1000; Cell Signaling; 2535), AMPK (1:1000; Cell Signaling 2793), phospho-ACC (1:1000; Cell Signaling; 3661), ACC (1:10,000; ProteinTech; 67373-1) and species-specific HRP-conjugated secondary antibodies (Vector Laboratories).

### RNA-seq

Sequencing was performed by the Advanced Genomics Core at University of Michigan Medical School. 250 human islets per donor were handpicked and washed twice with 1X PBS. 250,000 cells from differentiation stage 0 through 6 were washed twice with 1X PBS. Total RNA was isolated from human islet and stem cell samples and DNase treated using commercially available kits (Omega Biotek and Ambion, respectively). Libraries were constructed and subsequently subjected to 151 bp paired-end cycles on the NovaSeq-6000 platform (Illumina). FastQC (v0.11.8) was used to ensure the quality of data. Reads were mapped to the reference genome GRCm38 (ENSEMBL), using STAR (v2.6.1b) and assigned count estimates to genes with RSEM (v1.3.1). Alignment options followed ENCODE standards for RNA-seq. FastQC was used in an additional post-alignment step to ensure that only high-quality data were used for expression quantitation and differential expression. Data were pre-filtered to remove genes with 0 counts in all samples. Differential gene expression analysis was performed using DESeq2, using a negative binomial generalized linear model (thresholds: log2(fold change) >0.5 or <-0.5, FDR (P_adj_) <0.05). Plots were generated using variations of DESeq2 plotting functions and other packages with Genialis (Boston, MA, USA).

### Pathway analysis of differentially expressed genes

Differentially expressed genes for each cell-type were identified using DESeq2 packages with Genialis (Boston, MA, USA). The Kyoto Encyclopedia of Genes and Genomes (KEGG), gene ontology (GO), and WikiPathways databases via both Enrichr and STRING databases were used to determine association with particular biological processes, diseases, and molecular functions. The top pathways with an FDR-adjusted *p*-value < 5% were summarized in the results.

### Transcription factor analysis of differentially expressed genes

Differentially expressed genes for each cell-type were identified using DESeq2 packages with Genialis (Boston, MA, USA). ChIP Enrichment Analysis (ChEA) and TF Perturbations Followed by Expression databases via Enrichr were used to determine association with particular biological processes, diseases, and molecular functions. The top pathways with an FDR-adjusted *p*-value < 5% were summarized in the results.

### Removal of batch effects between differentiations

Bias in gene expression values was removed using a tool called ComBat-seq, using protocol selection as batch effect and treatment selection as group effect. ComBat-seq adjusted batch effects in RNA-seq count data. Normalized RNA-seq counts were uploaded into new collections on the Genialis Expressions. The counts were adjusted by mapping the quantiles of the empirical distributions of data to the batch-free distributions^103^. Batch effect adjustment was done using the sva (v3.48.0) R package^104^. The ComBat-Seq method adjusted raw gene counts for differentiation protocol while retaining biological signal from treatment. Batch effect corrected raw counts were TPM normalized using rnanorm (v1.5.1) Python package (https://github.com/genialis/RNAnorm).

### Analysis of publicly-available scRNA-seq and scATAC-seq^28^

Pseudo-bulk analyses were conducted by comparison of log2 (fold change) between SC-β cells compared to primary β cells. For mapping the SC-β cell differentiation trajectory, we downloaded preprocessed multi-omics (paired scRNA-seq and scATAC-seq) data from SC islets and performed quality control on each modality and sample individually and retained only barcodes that passed quality filtering in both modalities. We embedded the cells using 100 highly variable genes based on the transcriptomic modality only with phate^104^ and clustered the cells based on the first 2 phate components. Based on marker genes, we identified trajectories from immature to mature endocrine cell types and extracted the β cell lineage and binned cells along the first phate component. For identification of primary human β cells, we downloaded preprocessed multi-omics (paired scRNA-seq and scATAC-seq) data from primary human islets and performed quality control on each modality and sample individually and retained only barcodes that passed quality filtering in both modalities. We embedded the cells using UMAP based on the first 20 Harmony^105^ corrected principal components calculated from 2000 highly variable genes based on the transcriptomic modality. We clustered the cells and based on marker genes, retained only primary human β cells, which we concatenated with the cells extracted from the SC-β cell differentiation trajectory. For estimating motif activity, we called peaks based on the ATAC-seq modality with MACS2^106^ using the cells extracted from the SC-β cell differentiation trajectory and counted fragments in peaks for both primary and SC-β cells using Signac^107^. We identified motifs using the JASPAR2024 database^108^, which we used as input for chromVAR^51^.

### Transmission electron microscopy

Human islets and SC-islets were handpicked, fixed in 3% glutaraldehyde and 3% paraformaldehyde in 0.1 M Cacodylate buffer (CB; pH 7.2) at room temperature for 5 min and then stored at 4 °C. Islets were then washed with PBS and centrifuged at 2000 rpm for 2 min. Pre-warmed 2% agarose solution was then carefully added to islet pellets, centrifuged and allowed to cool at 4 °C for 30 min. Samples were then subjected to osmification in 1.5% K4Fe(CN)6 + 2% OsO4 in 0.1 CB for 1 h, dehydrated by serial washes in EtOH (30%, 50%, 70%, 80%, 90%, 95% and 100%) and embedded in Spurr’s resin by polymerization at 60 °C for 24 h. Polymerized resins were then sectioned at 90 nm thickness using a Leica EM UC7 ultramicrotome and imaged at 70 kV using a JEOL 1400 TEM equipped with an AMT CMOS imaging system. Mitochondrial structures (Aspect Ratio, Perimeter, Circularity) were analyzed and quantified by ImageJ.

### Immunofluorescence of cryosections

SC-islets and scaffolds containing SC-islets were picked/excised and fixed in 4% paraformaldehyde for 4 h at 4 °C. SC-islets were washed with PBS and centrifuged at 2000 rpm for 2 min. SC-islets were incubated in methylene blue at room temperature for 20 min and then washed 2x with PBS. Excised scaffolds were washed 3x with PBS for 10 min. SC-islets and scaffolds were stored in 30% sucrose in PBS for 16 h at 4 °C. SC-islets and scaffolds were embedded in OCT and frozen at −80 °C and cryosectioned. SC-islet sections were rinsed with 75% EtOH for 5 min at RT to remove methylene blue. SC-islet and scaffold sections were washed 3x with PBS, permeabilized with 0.2% Triton X-100 in PBS for 15 min, washed 3x with PBS and blocked at RT for 1 h with 5% donkey serum in PBT. Primary antibodies were added for 16 h at 4 °C. Primary antibodies raised against C-peptide (1:50; DSHB, Catalog#GN-ID4), Glucagon (1:2000; SantaCruz, Catalog# sc-13091), Glucagon (1:500; Cell Signaling, Catalog#2760), Somatostatin 28 (1:500; Abcam, Catalog#ab111912), Human antigen clone 235-1 (Millipore Sigma; Catalog#MAB1281), OXPHOS (1:100; Abcam, Catalog#ab110411), PDX1 (1:1000; Abcam, Catalog#ab47308), NKX6.1 (1:500; Abcam, Catalog#ab221549), and Ki-67 (1:250, Thermo, Catalog#PA5-19462) were used. Sections were then washed 3x with PBS and incubated for 2 h at room temperature with species-specific Dylight405, Cy2, Cy3, and Cy5-conjugated secondary antibodies (Jackson Immunoresearch). Nuclear labelling was performed using DAPI (Invitrogen). Images were captured under 20X and 60X oil immersion magnification on an Olympus IX81 microscope. Immunostained islets were captured with Z-stack images and subjected to deconvolution (CellSens; Olympus). Hormone-positive cells were quantified as hormone-positive cell/total DAPI-positive (*in vitro*) or Human Ag-positive (post-transplantation) cells. Ki-67-positive cells were quantified as Ki-67-positive/total C-peptide-positive cells. Quantification of hormone-positive and Ki-67-positive cells was carried out with blinded samples, and samples were reidentified after quantification to reduce bias. Quantitative 3-D assessments of mitochondrial morphology and network were performed on ImageJ using Mitochondria Analyzer plugin^102,109^.

### Statistics

In all figures, data are presented as mean ± SEM, and error bars denote SEM, unless otherwise noted in the legends. Statistical comparisons were performed using unpaired two-tailed Student’s *t*-tests, one-way or two-way ANOVA, followed by Tukey’s or Sidak’s post-hoc test for multiple comparisons, as appropriate (GraphPad Prism and R). A *P* value < 0.05 was considered significant. For transcriptomic profiling, a FDR-corrected *P* value <0.05 were considered significant (see RNA-seq of SC-islets and human islets above). Outlier tests (ROUT method^110^) were routinely performed in GraphPad Prism, and all results were included in analysis unless designated by the ROUT method.

## Data Availability

RNA-seq data have been deposited at GEO (GSE270220). Any additional information required to reanalyze the data reported in this paper is available from the lead author on request.

## Acknowledgements

S.A.S. acknowledges support from the JDRF (CDA-2016-189, COE-2019-861, SRA-2023-1392), the NIH (R01 DK108921, R01 DK135032, R01 DK135268, R01 DK136671, R01 DK127270, U01 DK127747, P30 DK020572), the Department of Veterans Affairs (I01 BX004444), the Brehm family, and the Anthony family. A.C.L. was supported by the NIH (T32-GM832230). J.L.K. was supported by the National Science Foundation Graduate Research Fellowship (DGE-2241144). G.L.P. was supported by the American Diabetes Association (19-PDF-063). E.L-D. was supported by the NIH (T32-AI007413, T32-AG000114). B.H-K. was supported by the NIH (T32-AI007413, T32-GM145304, F31-DK138544). E.M.W. was supported by the NIH (K01 DK133533). J.G.S.M was supported by the Novo Nordisk Foundation (NNF21OC0068929). S.P. was supported by the JDRF (COE-2019-861) and the NIH (P30DK089503). The JDRF Career Development Award to S.A.S. is partly supported by the Danish Diabetes Academy and the Novo Nordisk Foundation. Next generation sequencing was carried out in the Advanced Genomics Core at the University of Michigan. Human pancreatic islets and/or other resources were provided by the NIDDK-funded Integrated Islet Distribution Program (IIDP) (RRID:SCR _014387) at City of Hope, NIH Grant # U24DK098085. Human islets were also provided by the Alberta Diabetes Institute IsletCore at the University of Alberta in Edmonton (http://www.bcell.org/adi-isletcore.html) with the assistance of the Human Organ Procurement and Exchange (HOPE) program, Trillium Gift of Life Network (TGLN), and other Canadian organ procurement organizations. Islet isolation was approved by the Human Research Ethics Board at the University of Alberta (Pro00013094). All donors’ families gave informed consent for the use of pancreatic tissue in research. We thank Drs. K. Claiborn, L. Satin, R. Kennedy, D. Lorberbaum, S. Nowinski, and members of the Soleimanpour laboratory for helpful advice.

## Author Contributions

A.C.L. conceived, designed, and performed experiments, interpreted results, drafted, and reviewed the manuscript. E.M.W., J.K., K.C., E.B., A.M.S., J.Z., G.L.P, E.L-D., B.H-K., R.K.D., J.L., Y.W., E.C.R., M.A., V.S., J.P.P., and L.M. designed and performed experiments and interpreted results. T.J.H., M.M.C., J.G.S.M., S.P., and L.D.S. designed studies, interpreted results, and reviewed the manuscript. S.A.S. conceived and designed the studies, interpreted results, drafted, edited, and reviewed the manuscript.

## Conflict of interest statement

S.A.S has received grant funding from Ono Pharmaceutical Co., Ltd. and is a consultant for Novo Nordisk.

## Notes

### Summary of Updates

New and revised studies shared following external peer-review

