## Supplementary Information for "Limitations in PPAR⍺-dependent mitochondrial programming restrain the differentiation of human stem cell-derived β cells"

**Supplementary Table 1. Clinical characteristics of human islet donors**

| Unique Islet Prep Identifier | Islet Isolation Center | Age | BMI | Sex | Cause of Death |
|---|---|---|---|---|---|
| SAMN15770453 | IIDP | 48 | 30.9 | Female | Cerebrovascular/stroke |
| HP-20228-01 | Prodo Labs | 36 | 25.8 | Female | Anoxic event |
| HP-20234-01 | Prodo Labs | 58 | 26.9 | Female | Stroke |
| SAMN15877725 | IIDP | 31 | 27.4 | Male | Head trauma |
| HP-21213-01 | Prodo Labs | 62 | 23.0 | Female | Stroke |
| HP-21216-01 | Prodo Labs | 54 | 31.8 | Female | Stroke |
| HP-21227-01 | Prodo Labs | 58 | 31.8 | Male | Heart attack |
| HP-21232-01 | Prodo Labs | 52 | 29.6 | Male | Stroke |
| SAMN20923891 | IIDP | 45 | 50.6 | Male | Head trauma |
| HP-21244-01 | Prodo Labs | 59 | 20.7 | Female | Stroke |
| SAMN31040039 | IIDP | 28 | 27.8 | Male | Head trauma |
| HP-22272-01 | Prodo Labs | 58 | 29.0 | Male | Stroke |
| HP-22278-01 | Prodo Labs | 37 | 29.1 | Male | Anoxic event |
| HP-22294-02 | Prodo Labs | 36 | 24.4 | Male | Head trauma |
| R461 | ADI Islet Core, Alberta | 61 | 31.6 | Male | Neurological |
| HP-23046-01 | Prodo Labs | 60 | 32.9 | Male | Stroke |
| HP-23098-01 | Prodo Labs | 42 | 29.3 | Female | Stroke |
| SAMN36510137 | IIDP | 63 | 32.1 | Male | Anoxia |
| HP-23200-01 | Prodo Labs | 57 | 30.8 | Male | Anoxic event |
| SAMN36823227 | IIDP | 41 | 38.4 | Female | Cerebrovascular/stroke |
| HP-23219-01 | Prodo Labs | 59 | 24.5 | Male | Stroke |
| HP-23270-01 | Prodo Labs | 34 | 27.0 | Male | Stroke |
| HP-23258-01 | Prodo Labs | 39 | 26.9 | Male | Stroke |

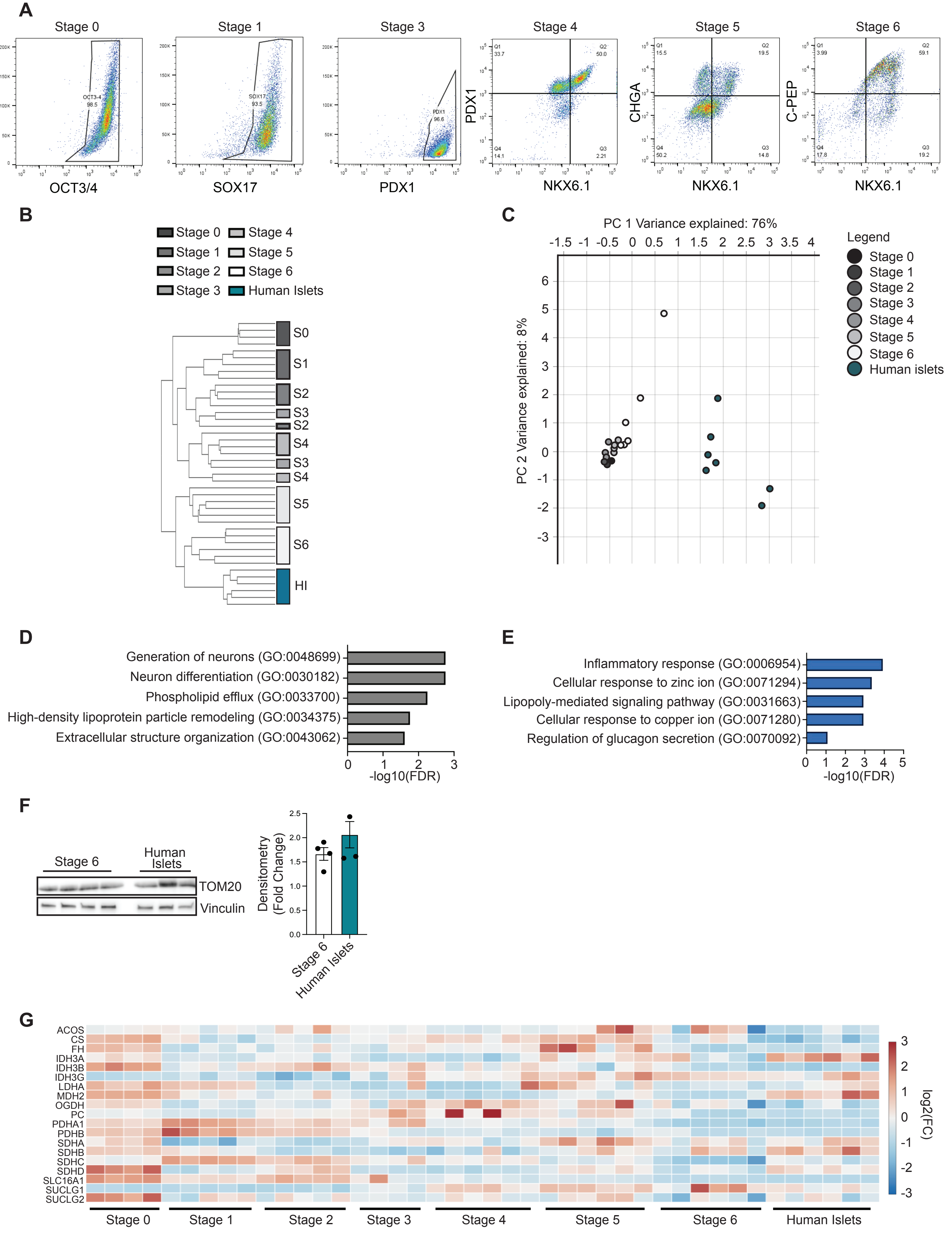
A
Stage 0
Stage 1
Stage 3
Stage 4
Stage 5
Stage 6
OCT3-4 98.5
SOX17 93.5
PDX1 96.6
OCT3/4
SOX17
PDX1
PDX1
CHGA
C-PEP
NKX6.1
Q1 33.7
Q2 50.0
Q4 14.1
Q3 2.21
Q1 15.5
Q2 19.5
Q4 50.2
Q3 14.8
Q1 3.99
Q2 59.1
Q4 17.8
Q3 19.2
B
Stage 0
Stage 1
Stage 2
Stage 3
Stage 4
Stage 5
Stage 6
Human Islets
S0
S1
S2
S3
S2
S4
S3
S4
S5
S6
HI
C
PC 1 Variance explained: 76%
PC 2 Variance explained: 8%
Legend
Stage 0
Stage 1
Stage 2
Stage 3
Stage 4
Stage 5
Stage 6
Human islets
D
Generation of neurons (GO:0048699)
Neuron differentiation (GO:0030182)
Phospholipid efflux (GO:0033700)
High-density lipoprotein particle remodeling (GO:0034375)
Extracellular structure organization (GO:0043062)
-log10(FDR)
E
Inflammatory response (GO:0006954)
Cellular response to zinc ion (GO:0071294)
Lipopoly-mediated signaling pathway (GO:0031663)
Cellular response to copper ion (GO:0071280)
Regulation of glucagon secretion (GO:0070092)
-log10(FDR)
F
Stage 6
Human Islets
TOM20
Vinculin
Densitometry (Fold Change)
Stage 6
Human Islets
G
ACOS
CS
FH
IDH3A
IDH3B
IDH3G
LDHA
MDH2
OGDH
PC
PDHA1
PDHB
SDHA
SDHB
SDHC
SDHD
SLC16A1
SUCLG1
SUCLG2
log2(FC)
Stage 0
Stage 1
Stage 2
Stage 3
Stage 4
Stage 5
Stage 6
Human Islets

**Figure S1. Differentiation of human embryonic stem cells generates transcriptionally immature SC-islets.** (A) Representative flow cytometry plots of stage specific markers from designated differentiation stages. (*n* = 3 differentiations/group). (B) Hierarchical clustering generated from RNA-seq for differentiation stages from the H1 cell line as well as human islet samples (*n* = 4-6 differentiations or independent human islet donors/group). (C) PCA plot generated from RNAseq for differentiation stages as well as human islet samples (*n* = 4-6 differentiations or independent human islet donors/group). Selected terms of Enrichr analysis of differentially expressed genes from stage 6 SC-islets *vs*. human islets analyzed with the GO Biological Process library for all (D) upregulated and (E) downregulated genes (*n* = 6 differentiations or independent human islet donors/group). (F) Left - Western blot (WB) of TOM20 protein expression for stage 6 SC-islets and human islet donors (*n* = 4 differentiations or 3 independent human islet donors/group). Each lane represents an independent stage 6 SC-islet differentiation or human islet donor. Vinculin serves as a loading control. Quantification (right) of TOM20 expression (normalized to Vinculin expression) via densitometry of the Western blots (*n* = 3-4/group). (G) Differential RNA expression heatmap, displayed as integrated mean $\log_2$ fold change (FC), from RNA-seq data of genes related to TCA cycle enzymes and lactate metabolism/transport at each differentiation stage and human islets (*n* = 4-6 differentiations or independent human islet donors/group).

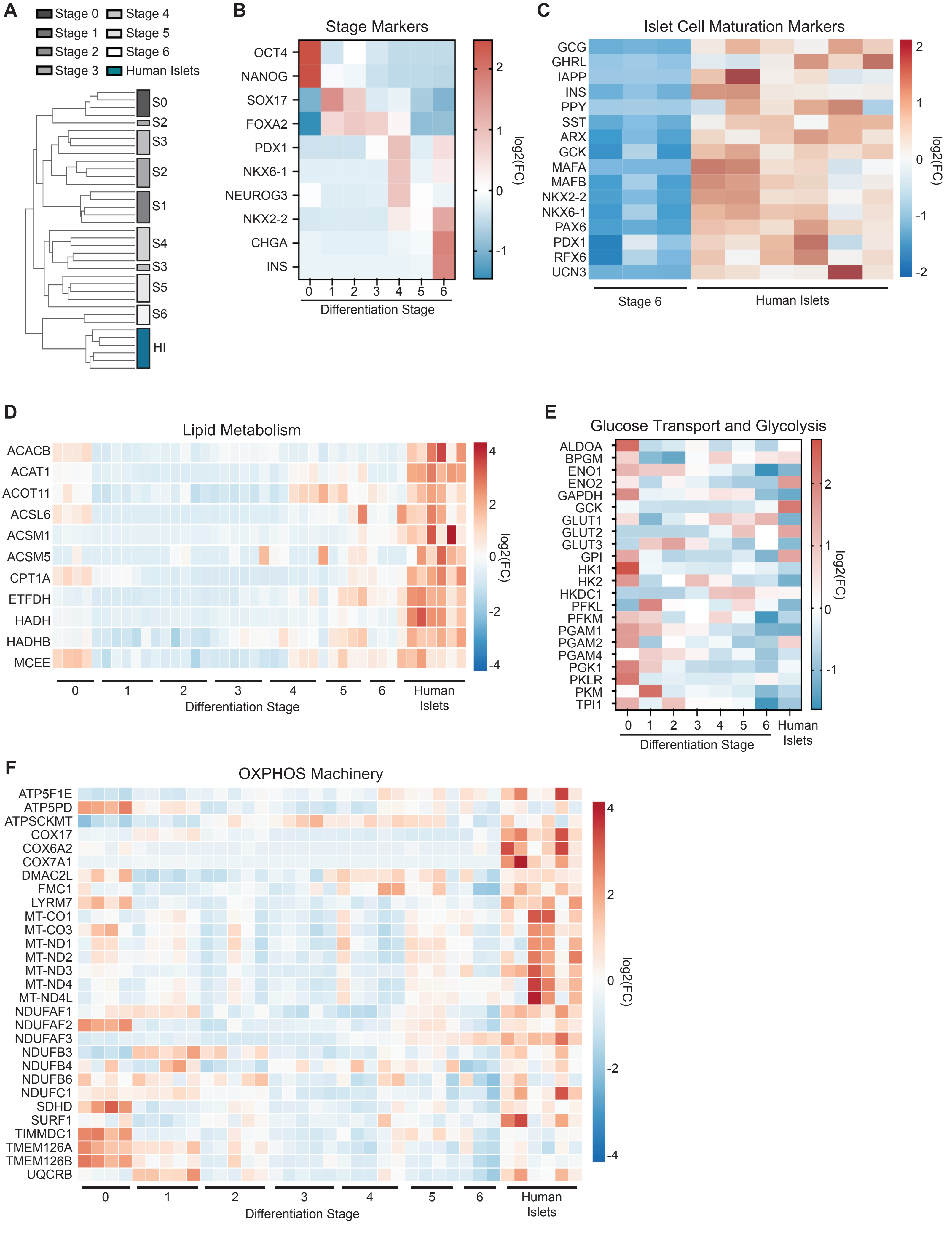

A
Stage 0
Stage 1
Stage 2
Stage 3
Stage 4
Stage 5
Stage 6
Human Islets
S0
S2
S3
S2
S1
S4
S3
S5
S6
HI
B
Stage Markers
OCT4
NANOG
SOX17
FOXA2
PDX1
NKX6-1
NEUROG3
NKX2-2
CHGA
INS
0 1 2 3 4 5 6
Differentiation Stage
log2(FC)
C
Islet Cell Maturation Markers
GCG
GHRL
IAPP
INS
PPY
SST
ARX
GCK
MAFA
MAFB
NKX2-2
NKX6-1
PAX6
PDX1
RFX6
UCN3
Stage 6
Human Islets
log2(FC)
D
Lipid Metabolism
ACACB
ACAT1
ACOT11
ACSL6
ACSM1
ACSM5
CPT1A
ETFDH
HADH
HADHB
MCEE
0 1 2 3 4 5 6 Human Islets
Differentiation Stage
log2(FC)
E
Glucose Transport and Glycolysis
ALDOA
BPGM
ENO1
ENO2
GAPDH
GCK
GLUT1
GLUT2
GLUT3
GPI
HK1
HK2
HKDC1
PFKL
PFKM
PGAM1
PGAM2
PGAM4
PGK1
PKLR
PKM
TPI1
0 1 2 3 4 5 6 Human Islets
Differentiation Stage
log2(FC)
F
OXPHOS Machinery
ATP5F1E
ATP5PD
ATPSCKMT
COX17
COX6A2
COX7A1
DMAC2L
FMC1
LYRM7
MT-CO1
MT-CO3
MT-ND1
MT-ND2
MT-ND3
MT-ND4
MT-ND4L
NDUFAF1
NDUFAF2
NDUFAF3
NDUFB3
NDUFB4
NDUFB6
NDUFC1
SDHD
SURF1
TIMMDC1
TMEM126A
TMEM126B
UQCRB
0 1 2 3 4 5 6 Human Islets
Differentiation Stage
log2(FC)

**Figure S2. Differentiation of HUES8 human embryonic stem cells generates SC-islets with transcriptional signatures of reduced mitochondrial oxidative and lipid metabolism.** (A) Hierarchical clustering generated from RNAseq for differentiation stages from the HUES8 cell line and human islet samples (*n* = 4-6 differentiations or independent human islet donors/group). (B) Differential RNA expression heatmap, displayed as integrated mean $\log_2$ fold change (FC), from RNAseq of representative stage marker expression at each differentiation stage from the HUES8 cell line (*n* = 4-5 differentiations/group). (C) Differential RNA expression heatmap of genes encoding islet cell maturation markers and hormones in HUES8 stage 6 SC-islets (*n* = 3 differentiations/group) and non-diabetic human islet donors (*n* = 6 independent human islet donors/group). FC is relative to the average across all individual samples. (D) Differential RNA expression heatmaps from RNA-seq data demonstrating expression of expression of lipid metabolism genes in HUES8 stage 0-6 cells and human islets (*n* = 3-5 differentiations or independent human islet donors/group). FC is relative to the average across all individual samples. (E) Differential RNA expression heatmap, displayed as integrated mean $\log_2$ fold change (FC), from RNA-seq data of genes related to glucose transport and glycolysis at each differentiation stage and human islets (*n* = 3-5 HUES8 differentiations or independent human islet donors/group). (F) Differential RNA expression heatmaps from RNA-seq data demonstrating expression of expression of OXPHOS machinery genes in HUES8 stage 0-6 cells and human islets (*n* = 3-5 differentiations or independent human islet donors/group). FC is relative to the average across all individual samples.

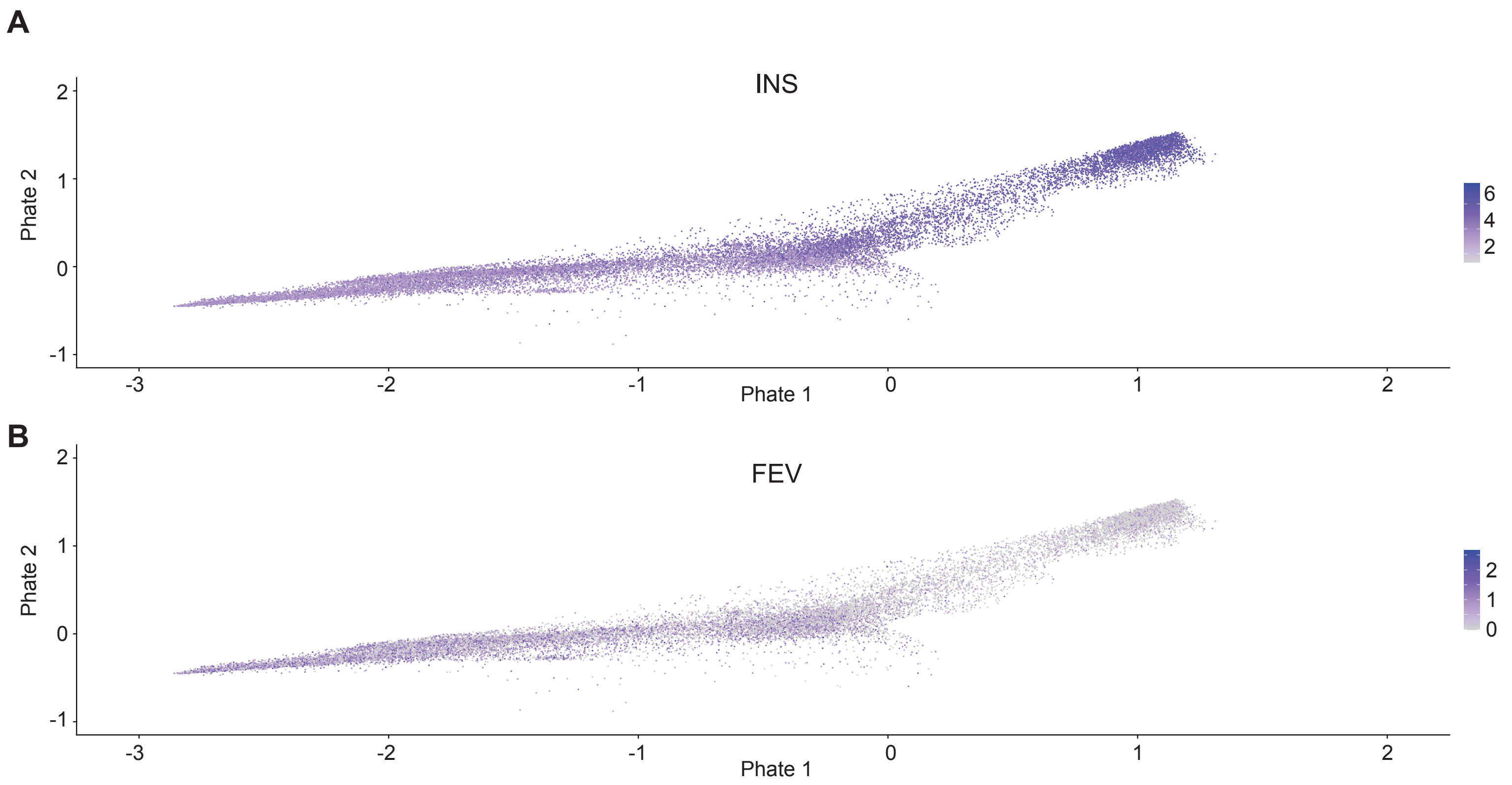
A
INS
Phate 2
Phate 1
6
4
2
B
FEV
Phate 2
Phate 1
2
1
0

**Figure S3**. **Stage 6 SC-β cells display increased insulin expression across SC-β cell differentiation trajectory.** UMAP projections of stage 6 SC-β cell differentiation trajectory and normalized expression of (A) *INS* and (B) *FEV* generated from re-analysis of scRNA-seq data from Augsornworawat *et al*[28].

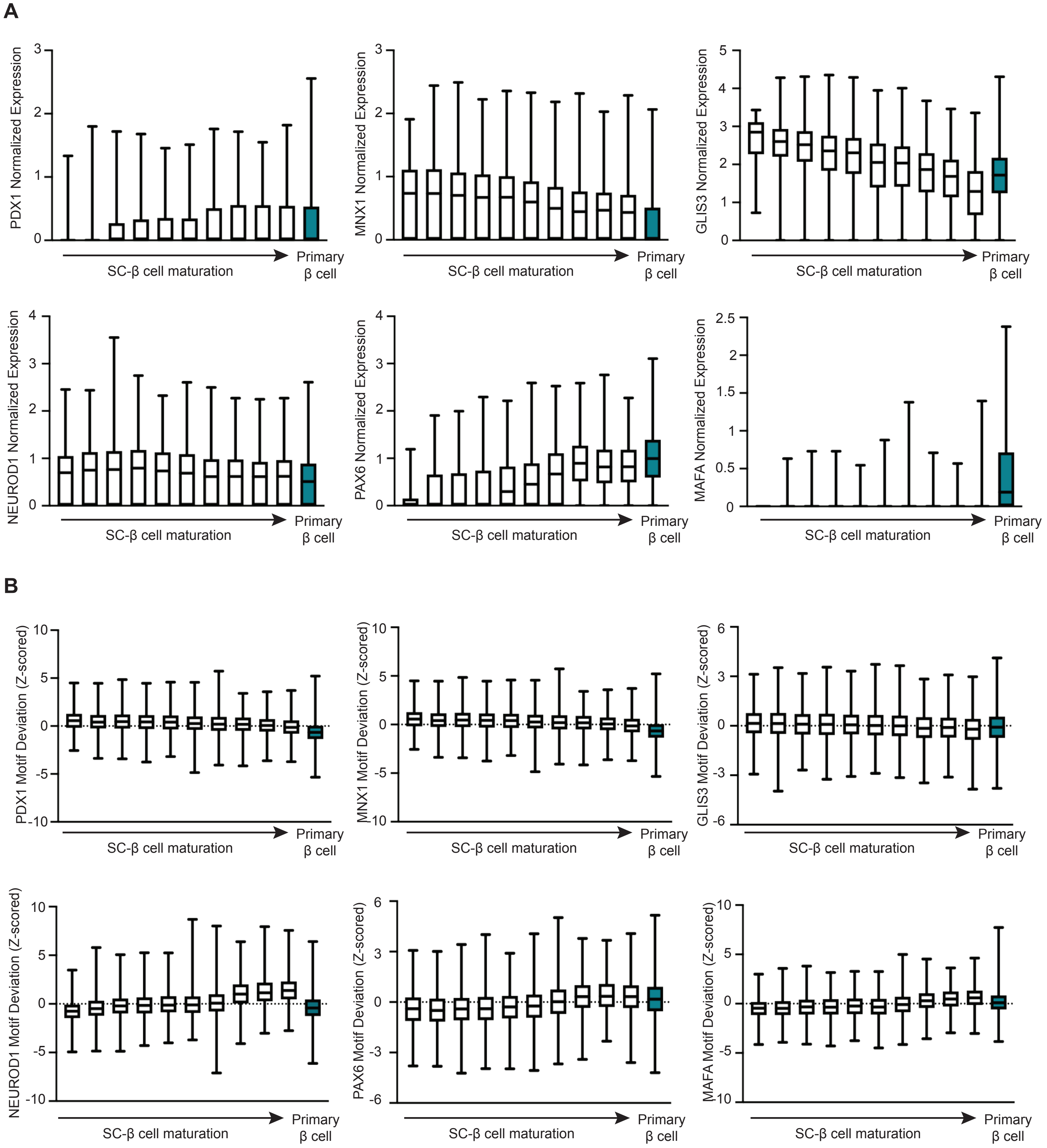

A
PDX1 Normalized Expression
0
1
2
3
SC-β cell maturation
Primary β cell
MNX1 Normalized Expression
0
1
2
3
GLIS3 Normalized Expression
0
1
2
3
4
5
NEUROD1 Normalized Expression
0
1
2
3
4
PAX6 Normalized Expression
0
1
2
3
4
MAFA Normalized Expression
0
0.5
1
1.5
2
2.5
B
PDX1 Motif Deviation (Z-scored)
-10
-5
0
5
10
MNX1 Motif Deviation (Z-scored)
-10
-5
0
5
10
GLIS3 Motif Deviation (Z-scored)
-6
-3
0
3
6
NEUROD1 Motif Deviation (Z-scored)
-10
-5
0
5
10
PAX6 Motif Deviation (Z-scored)
-6
-3
0
3
6
MAFA Motif Deviation (Z-scored)
-10
-5
0
5
10

**Figure S4. SC-β cells display enhanced maturation across SC-β cell differentiation trajectory.** Minimum to maximum box and whisker plots across SC-β cells differentiation trajectory and primary β cells generated from publicly-available scRNA-seq and scATAC-seq data[28] for normalized expression (A) and motif deviation z-scores (B), representing transcription factor motif activity, for the following transcription factors PDX1, MNX1, GLIS3, NEUROD1, PAX6, and MAFA (*n* = 3-4 differentiations or independent human islet donors/group).

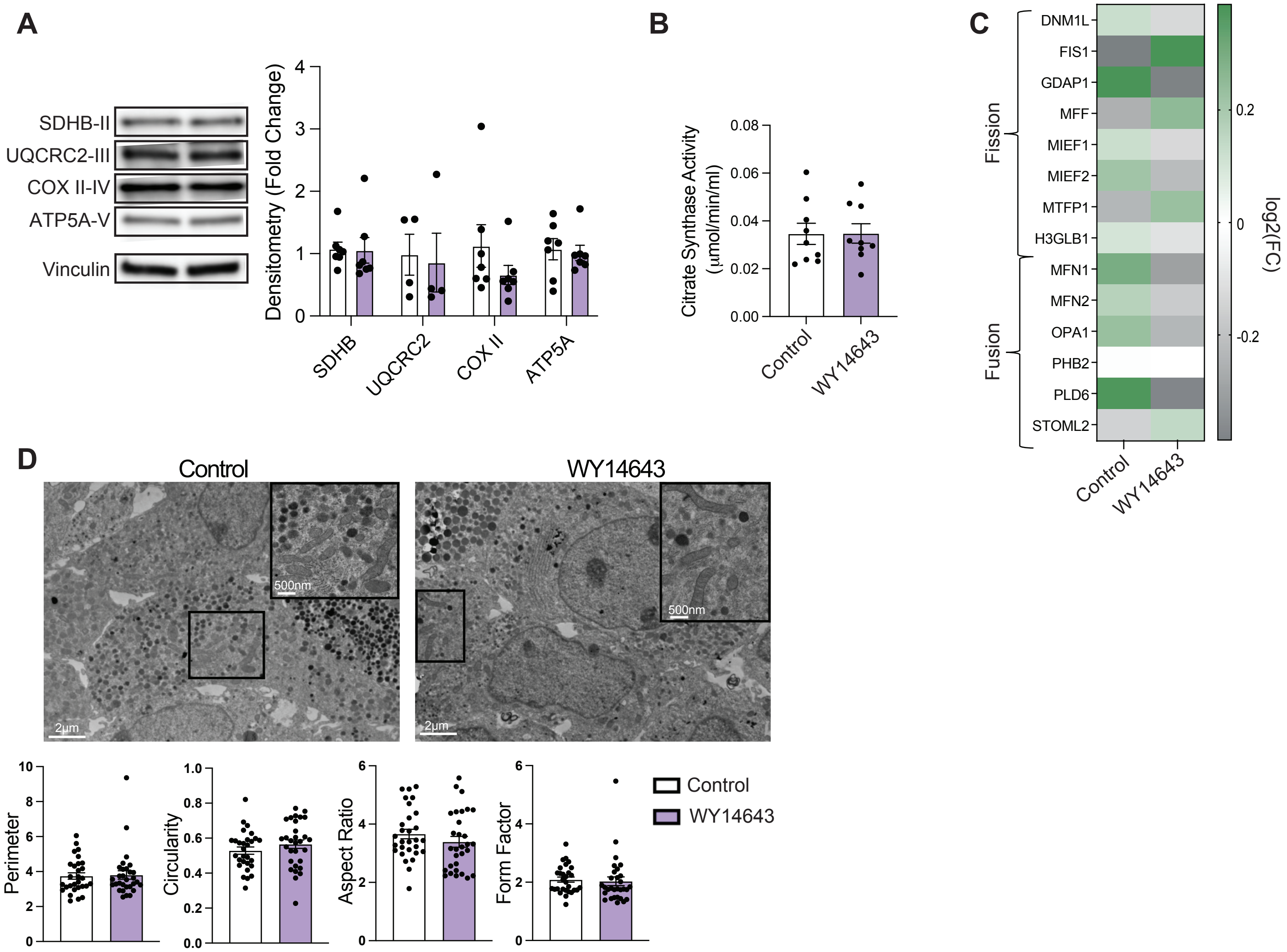

A
SDHB-II
UQCRC2-III
COX II-IV
ATP5A-V
Vinculin
Densitometry (Fold Change)
SDHB
UQCRC2
COX II
ATP5A
B
Citrate Synthase Activity (µmol/min/ml)
Control
WY14643
C
Fission
Fusion
DNM1L
FIS1
GDAP1
MFF
MIEF1
MIEF2
MTFP1
H3GLB1
MFN1
MFN2
OPA1
PHB2
PLD6
STOML2
log2(FC)
Control
WY14643
D
Control
WY14643
500nm
2µm
Perimeter
Circularity
Aspect Ratio
Form Factor
Control
WY14643

**Figure S5. WY14643 treatment does not alter mitochondrial mass or structure in SC-β cells.** (A) Representative Western blot analysis of expression of OXPHOS subunits, including SDHB (Complex II), UQCRC2 (Complex III), COXII (Complex IV), and ATP5A (Complex V) in vehicle control and WY14643-treated stage 6 SC-islets (left). Vinculin serves as a loading control. Quantification (right) of SDHB, UQCRC2, COXII, and ATP5A protein expression by densitometry, normalized for Vinculin expression. (*n* = 4-6 differentiations/group). (B) Quantification of citrate synthase activity normalized to total protein content in vehicle control and WY14643-treated stage 6 SC-islets (*n* = 9 differentiations/group). (C) Differential RNA expression heatmap (presented as integrated mean/group) of genes encoding mitochondrial fission and fusion regulators in vehicle control and WY14643-treated stage 6 SC-islets (*n* = 4/group). (D) Representative transmission electron microscopy (TEM) images of SC-β cells in vehicle control and WY14643-treated stage 6 SC-islets and quantification of mitochondria ultrastructure (right) (*n* = 3/group, ~150 β cell mitochondria scored/sample).

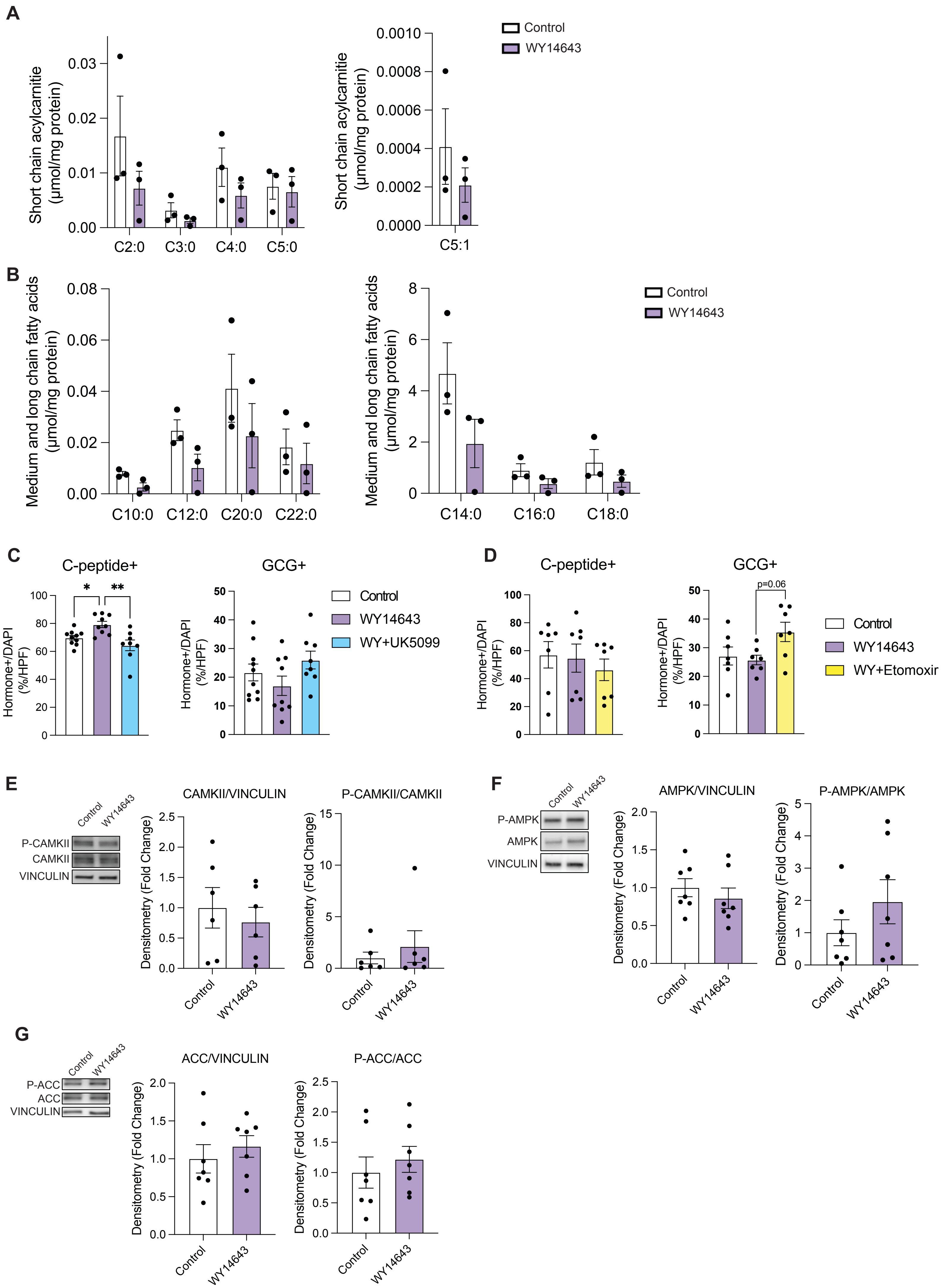
A
Short chain acylcarnitie (µmol/mg protein)
0.03
0.02
0.01
0.00
C2:0
C3:0
C4:0
C5:0
0.0010
0.0008
0.0006
0.0004
0.0002
0.0000
C5:1
Control
WY14643
B
Medium and long chain fatty acids (µmol/mg protein)
0.08
0.06
0.04
0.02
0.00
C10:0
C12:0
C20:0
C22:0
8
6
4
2
0
C14:0
C16:0
C18:0
Control
WY14643
C
C-peptide+
GCG+
Hormone+/DAPI (%/HPF)
*
**
Control
WY14643
WY+UK5099
D
C-peptide+
GCG+
p=0.06
Control
WY14643
WY+Etomoxir
E
Control
WY14643
P-CAMKII
CAMKII
VINCULIN
CAMKII/VINCULIN
P-CAMKII/CAMKII
Densitometry (Fold Change)
F
P-AMPK
AMPK
VINCULIN
AMPK/VINCULIN
P-AMPK/AMPK
G
P-ACC
ACC
VINCULIN
ACC/VINCULIN
P-ACC/ACC

**Figure S6. WY14643 treatment improves mitochondrial oxidative and fatty acid metabolism to promote SC-β cell differentiation.** Lipidomics profiling of (A) short-chain acylcarnitines or (B) medium- and long-chain fatty acids (normalized to protein content) in vehicle control and WY14643-treated stage 6 SC-islets (*n* = 3 differentiations/group). (C) Quantification of C-peptide-positive (left) or glucagon-positive cells (right) per high powered field (HPF) in stage 6 SC-islets following 7 days of treatment with vehicle control, WY14643 alone, or WY14643 + 4 μM UK5099 (*n* = 8-10 fields quantified/group, ~1000 cells quantified/differentiation and respective condition from 2 differentiations/group). *$P < 0.05$, **$P < 0.01$ by one-way ANOVA followed by Tukey's multiple comparisons test. (D) Quantification of C-peptide-positive (left) or glucagon-positive cells (right) per high powered field (HPF) in stage 6 SC-islets following 7 days of treatment with vehicle control, WY14643 alone, or WY14643 + 25 μM etomoxir (*n* = 7-8 fields quantified/group, ~1000 cells quantified/differentiation and respective condition from 2 differentiations/group). (E) Representative Western blot analysis (left) with densitometry (right) of expression of total CAMKII or phosphorylated CAMKII (P-CAMKII) in vehicle control and WY14643-treated stage 6 SC-islets (left). Vinculin serves as an additional loading control. (*n* = 6 differentiations/group). (F) Representative Western blot analysis (left) with densitometry (right) of expression of total AMPK or phosphorylated AMPK (P-AMPK) in vehicle control and WY14643-treated stage 6 SC-islets (left). Vinculin serves as an additional loading control. (*n* = 7 differentiations/group). (G) Representative Western blot analysis (left) with densitometry (right) of expression of total ACC or phosphorylated ACC (P-ACC) in vehicle control and WY14643-treated stage 6 SC-islets (left). Vinculin serves as an additional loading control. (*n* = 7 differentiations/group).

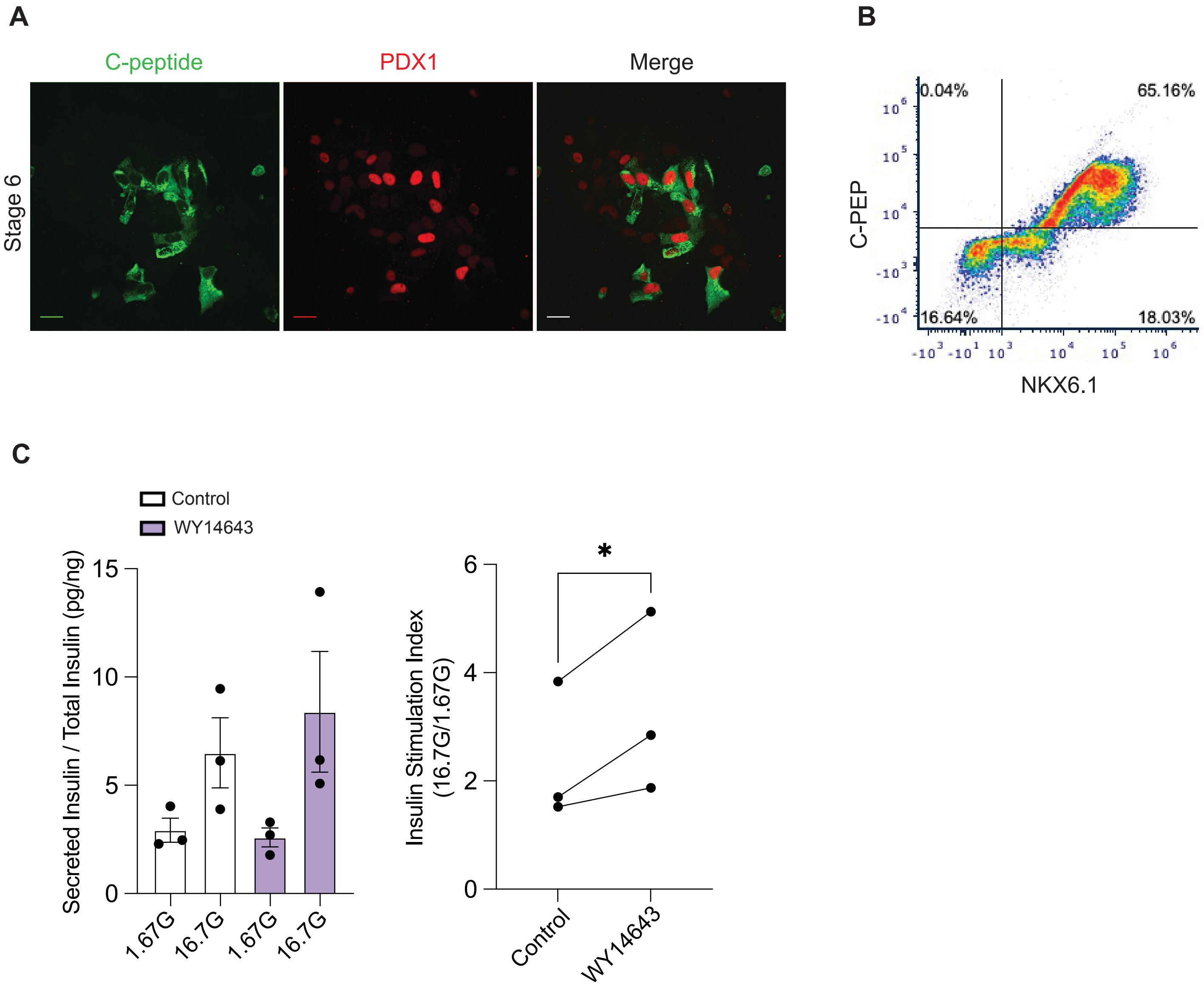

A
C-peptide
PDX1
Merge
Stage 6
B
0.04%
65.16%
16.64%
18.03%
C-PEP
NKX6.1
C
Control
WY14643
Secreted Insulin / Total Insulin (pg/ng)
15
10
5
0
1.67G
16.7G
1.67G
16.7G
Insulin Stimulation Index (16.7G/1.67G)
6
4
2
0
*
Control
WY14643

**Figure S7. WY14643 treatment improves insulin secretion in stage 6 iPSC-islets derived from 19-9-11 cells.** (A) Immunofluorescence images at 60X magnification of stage 6 iPSC-islets for C-peptide (green) and PDX1 (red) (*n* = 3/group). Scale bars, 20 μm. (B) Representative flow cytometry plots from stage 6 iPSC-islets in culture. (*n* = 3/group). (C) Insulin secretion (left) with insulin stimulation index (right) following static incubations in 1.67 mM glucose (1.67G) and 16.7 mM glucose (16.7G), normalized to total insulin content, in vehicle control and WY14643-treated stage 6 iPSC-islets. Stage 6 iPSC-islets were treated from days 27-34 before insulin secretion assays (*n* = 3 differentiations/group). $^{*}P < 0.05$ by Student's *t*-test.
